# The impact of the rice production system (irrigated vs lowland) on root-associated microbiome from farmer’s fields in western Burkina Faso

**DOI:** 10.1101/2022.03.29.486073

**Authors:** Mariam Barro, Issa Wonni, Marie Simonin, Abalo Itolou Kassankogno, Agnieszka Klonowska, Lionel Moulin, Gilles Béna, Irénée Somda, Caroline Brunel, Charlotte Tollenaere

## Abstract

As a consequence of its potential applications for food safety, there is a growing interest in rice root-associated microbial communities, but some systems remain understudied. Here, we compare the assemblage of root-associated microbiota in rice sampled in 19 small farmer’s fields from irrigated and rainfed lowlands in western Burkina Faso, using an amplicon metabarcoding approach 16S (Prokaryotes, three plant sample per field) and ITS (fungi, one sample per field). In addition to the expected structure according to the root compartment (root vs. rhizosphere) and geographical zones, we show that the rice production system is a major driver of microbiome structure, both for prokaryotes and fungi. In irrigated systems, we found a higher diversity of prokaryotic communities from rhizosphere and more complex co-occurrence networks, compared to rainfed lowlands. Core taxa were different between the two systems, and indicator species were identified: mostly within Bacillaceae and Bradyrhizobiaceae families in rainfed lowlands, and within Burkholderiaceae and Moraxellaceae in irrigated areas. Finally, phylotypes assigned to putative phytobeneficial and pathogen species were found. Mycorrhizal fungi Glomeromycetes abundance was higher in rainfed lowlands. Our results highlight deep microbiome differences induced by contrasted rice production systems that should consequently be considered for potential microbial engineering applications.

## Introduction

Soil and rhizosphere host megadiverse and dynamic communities of microorganisms that are crucial to the plants they associate with. Their role is particularly recognized for crops as the below-ground microbiota supply plants with nutrients and provide protection against pathogens (Singh *et al*. 2020; Chialva *et al*. 2021). Recent research suggests that root-associated microbes can improve plant tolerance to environmental stressors (Chialva *et al*. 2020), and modify phenology (Lu *et al*. 2018) and morphological traits (Senthil Kumar *et al*. 2018). Cultivated plants and their associated microbial communities are thus increasingly studied jointly, as holobionts (a concept reviewed by Vandenkoornhuyse *et al*. in 2015), because a deeper understanding of their interaction might help to develop microbial engineering applications for modern sustainable agricultural systems (Chialva *et al*. 2021). While much progress has been made, the mechanisms that control root-associated microbiome assembly (i.e., structure, composition and dynamics) remain difficult to disentangle (Brunel *et al*. 2020). Vertical transmission of the microorganisms (i.e., across plant generation) exists as well as the horizontal transmission (i.e., recruitment from the soil “seed bank”), with variable contributions of the seed and soil to the seedling microbiome (Rochefort *et al*. 2021; Walsh *et al*. 2021). Beyond the environmental drivers known to shape bulk soil communities (e.g., climate, soil properties, agricultural practices; see Vieira *et al*. 2020), the influence of the cultivated plant in structuring communities is more and more documented and now referred as the extended root phenotype (see de la Fuente Cantó *et al*. 2020). Indeed, while the root-associated micro-habitats (i.e., rhizosphere, rhizoplane, endosphere) modulate the intensity of assembly processes (Beattie, 2018), the plant identity (e.g., species, genotype, age, etc.) plays a major role in recruiting specific microbial taxa shared across multiple environmental conditions (Schweitzer *et al*. 2008).

Rice is the most important food crop in the world, grown in variable climatic conditions and representing the staple food of more than half of the world’s population, mostly in Asia, Africa and Latin America (Pandey *et al*. 2010). The major species cultivated worldwide is *Oryza sativa L.* (known as ‘Asian rice’). This is also true in Africa, where a second rice species was domesticated (*O. glaberrima,* referred to as ‘African rice’), but is much less grown because of lower yield (Linares, 2002). Given its importance for food security, and the impact of microbiota on plant productivity, it is not surprising that many recent studies used metabarcoding approaches to describe the microbiome of *O. sativa* (Kim & Lee, 2020), particularly its root-associated microbial communities (reviewed by Ding *et al*. in 2019). Rice has the particularity to be cultivated in flooded paddy soils over most growth stages, so that the rhizosphere is located in an oxic-anoxic interface (Ding *et al*. 2019). Microbial communities inhabiting rice roots are distinct from those found in other crops, with for example an enrichement in *Deltaproteobacteria*, *Euryarchaeota*, *Chytridiomycota* (Ding *et al*. 2019). On the other hand, like for other crops, their structuring is driven both by the host plant (in terms of root compartment / microhabitats, plant genotype) and its environment (geographical zone, bioclimate, soil properties, agricultural practices; Ding *et al*. 2019). In terms of microhabitats, differences between rhizophere and endosphere compartments are clear (see e.g. Edwards *et al*. 2015; Guo *et al*. 2021) and recent evidence shows differences in microbiota composition between root types and along root axes (Kawasaki *et al*. 2021). An effect of host genotype (subspecies/cultivars) were evidenced in some cases (Edwards *et al*. 2015; Alonso *et al*. 2020), but it is generally weak compared to other factors, or even absent (Edwards *et al*. 2018; Guo *et al*. 2021). Environmental factors, such as the geographical zone and agricultural practices, are important drivers of microbiota structuring as well. For example, Edwards *et al*. (2015) evidenced an effect of geographical location and cultivation practices, namely organic *vs* conventional farming. Other environmental factors, such as drought stress (Santos-Medellín *et al*. 2017), water management (Chialva *et al*. 2020), phosphorus (Long & Yao, 2020), were also shown to affect rice root-associated microbiota. In addition, bacterial and archaeal communities evolve during the vegetative phase of plant growth, then shift and stabilize compositionally at the transition to reproductive growth at flowering stage (Edwards *et al*. 2018). Vertical transmission through seeds seems quite weak or even absent (Guo *et al*. 2021). However, the rice microbiome has been poorly explored in African context in spite of **(1)** the importance to investigate a diversity of geographical areas (as a consequence of the effect of geographical zone, see above) and document the diversity of cultural practices, and **(2)** the growing importance of rice in Africa (recent surge in rice consumption with 8% yearly increase from 2009 to 2019, Soullier *et al*. 2020). An exception is the recent work by Kanasugi *et al*. (2020) that evidenced an effect of the region in structuring of rice microbiome described in six tropic savanna regions in Ghana. More generally, there is a lack of knowledge concerning crop-associated microbiota in the African continent, that results in a biased view of the microbial world associated with crops due to worldwide sampling repartition (Brunel *et al*. 2020; Hughes *et al*. 2021), and it is of particular concern due to the potential of microbial engineering for future agriculture.

Rice is grown around the world in a diversity of rice-growing systems, three of them being the most important (Rao *et al*. 2017). First, irrigated lowlands, with full water control, produce 75% of the global rice production. Second, rainfed lowlands (including flood prone), represents around 19% of the world’s rice production. Finally, rainfed upland rice, only possible under high rainfall, results in 4% of the global total rice production. In Burkina Faso, irrigated rice represents small areas (costly infrastructures representing less than 30% of harvested areas; CountrySTAT, 2020), but produces more than half of the national rice production (MAHRH, 2011). On the other hand, rainfed lowlands represent the majority of rice growing surfaces (67% between 1984 and 2009), but only 42% of the production as a consequence of lower yields compared to irrigated areas (MAHRH, 2011). Irrigated areas (IR) and rainfed lowlands (RL) host different agricultural practices in West Africa (Nonvide *et al*. 2018). In western Burkina Faso in particular, these contrasted practices have been documented, showing that the possibility to grow rice twice a year was restricted to irrigated areas, while direct sowing was performed only in rainfed lowlands, and that mineral fertilization was more frequent in irrigated areas (Barro *et al*. 2021). In terms of the host plant genetic diversity however, a study led in six locations from western Burkina Faso revealed few differences between irrigated areas and rainfed lowlands in general, except in one of the study sites, the rainfed lowland of Karfiguela, highly differentiated from the other five sites (Barro *et al*. 2021).

This study aims at describing rice root-associated microbial communities in farmer’s fields from western Burkina Faso. More specifically, we investigate whether rice roots from two contrasted rice growing systems (irrigated and rainfed lowland areas) host different microbial communities. To this aim, we collected rice root and rhizosphere samples in farmer’s fields from three geographical zones, each consisting of an irrigated rice-growing area and neighboring rainfed lowlands. A previous study performed in the same study sites documented more intensive agricultural practices in irrigated areas, compared to rainfed lowlands (Barro *et al*. 2021). Considering the effect of intensification on belowground biodiversity and microbial network complexity (Banerjee *et al*. 2019; Tamburini *et al*. 2020), we hypothesize an effect of the rice growing system on root-associated microbial communities diversity and complexity. If true, this may have consequences on plant health (Wei *et al*. 2015), particularly in this system where rice diseases were shown to circulate at higher levels in irrigated areas compared to rainfed lowlands (Barro *et al*. 2021).

## Material and methods

### Study sites in western Burkina Faso

The study sites are located in three geographic zones in western Burkina Faso, with maximum distance between each zone about 90 kilometers (Fig. 1a). Each zone comprises one irrigated area and the neighboring rainfed lowland, with maximum distance between the two rice growing systems of each zone being 7 kilometers (Fig. 1b). The climate consequently do not differ between rice growing systems within each zone, but average precipitation during the rice growing season (July to early December) differ between the three geographical zones (WorldClim 2 data; Fick & Hijmans, 2017; see Fig. S1).

**Figure 1.**
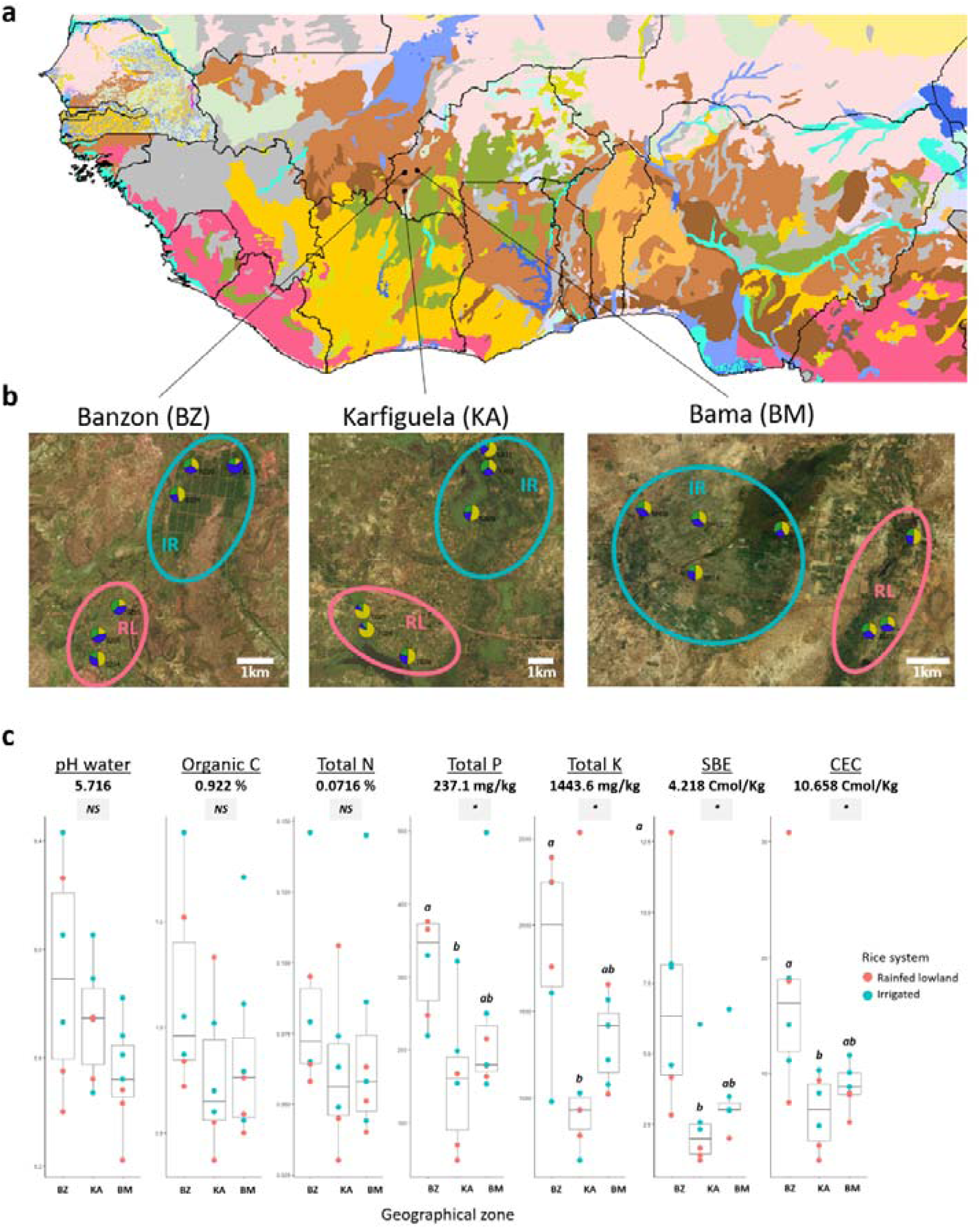
Location of the study sites and soil physico-chemical properties Location of the study sites in western Burkina Faso on the soil Harmonized Word Soil Database (HWSD) map of West Africa (https://webarchive.iiasa.ac.at/Research/LUC/External-World-soil-database/HTML/). The three geographical zones studied are in Lixisols (LX: soils with subsurface accumulation of low activity clays and high base saturation) Location of the field studied within each geographical zones. Soil texture, are indicated for each of the 19 studied fields, with pie charts representing relative proportions of sand (in yellow), silt (in green), and clay (in blue). Soil chemical properties estimated in each geographical zones, with colors representing the rice growing system (irrigated areas in blue and rainfed lowlands in red). Each point corresponds to one field studied. Averages over the 19 studied fields are indicated for each parameter, as well as the results of statistical tests for the geographical zone effect.

These six sites were studied from 2016 to 2019, with the characterization of agricultural practices and the follow-up of major rice diseases symptoms (Barro *et al*. 2021; see further details on the methodology and raw data at: https://doi.org/10.23708/8FDWIE). Rice genotyping data on samples collected in 2018 in these six sites are also available (see Barro *et al*. 2021).

### Rice root sampling

Within the six sites, we investigated a total of 19 fields, with three fields per site, except in Bama, the largest irrigated perimeter studied, where four fields were sampled (Fig. 1b). Each field studied corresponds to a square of approximately 25 meters on a side. Root and rhizosphere sampling was performed at rice maturation, between October, 16^th^ and December 3^rd^ 2018. We chose this developmental stage based on a previous study showing that the rice microbiome composition evolves during the growing season until a ‘mature’ microbiome at the flowering stage (Edwards *et al*. 2018). Within each field, we sampled three plants located on the square diagonal, with at least 5 meters distance. Sampling was performed with gloves and scissors (ethanol sanitized between two sampling) and involved nodal, bases and seminal roots (all sampled together). Roots were roughly shaken to remove non-adherent soil, and placed in 50 mL sterile tubes containing sterile Phosphate Buffered Saline (PBS) solution for a rapid (15s) rinse and then stored in another 50 mL sterile PBS-containing tube. We placed the tubes in a cooler and then at 4°C when back to the laboratory, on the same day.

### Soil physicochemical properties: data acquisition and analysis

The three geographical zones studied (Fig. 1a) are characterized by Lixisols (soils with subsurface accumulation of low activity clays and high base saturation) according to the Harmonized Word Soil Database (HWSD) map (FAO/IIASA/ISRIC/ISSCAS/JRC, 2012).

Soil sampling was performed on the same day as rice root sampling, and in three locations nearby sampled plants, using a 10 cm depth auger. Back from the field, sampled soil was dried in the shade at room temperature and stored until analysis.

INERA/GRN service performed the analyses of soil samples according to a standardized methodology. Briefly, soil physical properties were assessed by soil particle size distribution following Bouyoucos (1962), pH was estimated according to AFNOR (1981), total organic carbon with the Walkley & Black (1934)’s method, total concentrations of nitrogen with Kjeldhal method Hillebrand *et al*. (1953), and finally phosphorus and potassium content as described respectively in Novozansky *et al*. (1983) and Walinga *et al*. (1989). Then, cation exchange capacity (CEC), a measure of fertility, nutrient retention capacity, and the capacity to protect groundwater from cation contamination, was estimated, as well as sorptive bioaccessibility extraction (SBE), that relates to the environmental mobility, partitioning and toxicity of soil pollutants, following Metson (1956). Soil data are publicly available on the IRD Dataverse : https://doi.org/10.23708/LZ8A5B.

### Rice root conditioning, DNA extraction and sequencing

Less than 24h after sampling, root samples (rice roots including rhizosphere) stored at 4°C were processed. In order to separate the different root compartments, the tubes were vortexed vigorously one minute and then, roots were removed from the PBS solution using sterile forceps. The remaining PBS solution was considered as the ‘rhizosphere’ compartment. Roots were then surface-sterilized with 70% alcohol (30s), 1% bleach (30s) and finally rinsed three times in sterile water. We considered these surface-sterilized roots as the ‘root’ compartment, which comprises both endosphere microorganisms as well as persistent DNA from the rhizoplane. DNA extraction from the rhizosphere and root samples were performed on the same day as the process of compartment separation.

For DNA extractions, 0.25g of root samples (crushed in liquid nitrogen beforehand) and 0.25g of rhizosphere were extracted using the DNeasy PowerSoil Kit (Qiagen, Hilden, Germany), following manufacturer’s recommendations. DNA quality and quantity was verified using a NanoDrop ND-1000 spectrophotometer. PCR amplification, library and MiSeq Illumina sequencing were performed by Macrogen (Seoul, South Korea) using primers 341F (16S V3F, 5’-CCTACGGGNGGCWGCAG-3’) and 785R (16S V4R, 5’-GACTACHVGGGTATCTAATCC-3’) to amplify the V3 -V4 regions of the 16S rRNA gene, and using primers ITS1f: CTTGGTCATTTAGAGGAAGTAA and ITS2: GCTGCGTTCTTCATCGATGC to amplify the internal transcribed spacer 1 (ITS1) region. We specifically focused on the factors structuring prokaryotic communities (16S sequencing), but were also interested to see whether similar tendencies hold for fungal communities (ITS sequencing). We consequently chose the following approach: sequencing was performed for each sampled plant (19 fields * 3 plants = 57 samples per compartment) for 16S sequencing and for a composite sample (3 plant samples were pooled to result in one sample per field, so that the total number of samples per compartment is 19) for the ITS marker. Negative controls (three for 16S and one for ITS) were sequenced to remove potential contaminants.

Sequence data are retrievable from NCBI (National Center for Biotechnology Information) under the Bioproject ID: PRJNA763095.

### Bioinformatic analyses of obtained sequences

All bioinformatics and statistical analyses were performed in R software v 3.6.3 (R core Team, 2018) and the package *ggplot2* was used for the visualization.

Raw sequences were processed using a custom script from the *dada2* pipeline, which is designed to resolve exact biological sequences (ASVs for Amplicon Sequence Variants) from Illumina sequence data without sequence clustering (Callahan *et al*. 2016). Raw sequences were first demultiplexed by comparing index reads with a key, and paired sequences were trimmed. Sequences were dereplicated, and the unique sequence pairs were denoised using the *dada* function. Primers and adapters were screened and removed using a custom script with *cutadapt* (Martin, 2011). Next, paired-end sequences were merged, and chimeras were removed. Contaminants were identified and removed using negative controls and the *decontam* package. Rarefaction curves were drawn for each sample, using the *rarecurve* function of the *vegan* R package, and the rarefaction plateau was reached for all samples (Fig. S2). To account for differences in sequencing depths, samples were rarefied to 4236 and 7828 for 16S and ITS, respectively. Taxonomy assignments were determined against the UNITE 2021 (Abarenkov *et al*. 2021) and the SILVA SSU r138 (Quast *et al*. 2013) taxonomic databases for ITS and 16S, respectively using the *idtaxa* function from the *decipher* R package. Mitochondria and chloroplast sequences were then removed. ASVs not seen more than once in at least 2% of the samples were removed. We then obtained 2 116 969 (8260 ASVs) final sequences for 16S and 172 719 (566 ASVs) sequences for ITS. A figure showing the phyla relative abundances was constructed and presented according to the rice-growing system, site and root compartment.

Indices of α-diversity (observed richness and Shannon diversity index) were calculated using the *estimate_richness* function from the *phyloseq* package (McMurdie & Holmes, 2013). Members of the core microbiota were identified for 16S and ITS communities (including both rhizosphere and roots compartments) in each rice grown system using the prevalence threshold of 60%.

For 16S dataset only, we inferred co-occurrence networks using the *SpiecEasi* pipeline (Kurtz *et al*. 2015), independently for each rice growing system: rainfed lowland and irrigated areas. Networks were calculated for ASVs present in more than 15% of the samples using the method ’mb’ and setting the lambda.min.ratio to 1e-3 and nlambda to 50. We identified hub taxa, i.e. the potential keystone of the microbial network belonging to the most connected ASVs, based on their node parameters (method developed by Berry & Widder, 2014): a low betweenness centrality (lower quantile, < 0.9), and a high closeness centrality (higher quantile, > 0.75), transitivity (higher quantile, > 0.25) and degree (higher quantile, > 0.75). The node and network parameters were determined using the R package *igraph* (Csardi & Nepusz, 2006) and *qgraph* (Epskamp *et al*. 2012). Complete networks were further described by calculating the number of nodes and hubs, the network mean degree, mean closeness and betweenness centralities, the total number of edges, and the positive to negative edges ratio. Core taxa (prevalent in more than 60% of the samples) were also identified for each system.

The taxonomic affiliation of all ASVs was refined using nucleotide basic local alignment search tool (BLASTn) analyses on NCBI nr database. We screened the table of blast best hits of all 16S and ITS ASVs in order to search the genus or species names of a number of pathogen species listed a priori (Table S6), based on the reference book *Compendium of Rice Diseases* (Cartwright *et al*. 2018). We also searched within blast best hits for ITS ASVs assigned to the *Glomeromycetes* class, as well as 16S and ITS ASV assigned to a species name’s including “oryz”, as this may correspond to a specific interaction (pathogen or beneficial) with rice.

### Statistical analyses

We first analyzed soil physico-chemical parameters. To this purpose, PERMANOVA were performed on soil physical properties (texture, i.e. relative amount of sand, silt, and clay) on the one hand, and soil chemical properties (7 variables: pH, total carbon, total nitrogen, total phosphorus, total potassium, SBE, CEC) on the other hand. For both models, we included as explanatory factors the ‘geographical zone’ and ‘rice-growing system’, as well as their interaction, using *adonis2* function from the *vegan* R package (Oksanen *et al*. 2007), with 999 permutations. Posthoc tests were done using the *pairwiseAdonis* function (https://github.com/pmartinezarbizu/pairwiseAdonis). In addition, we performed Kruskal-Wallis non-parametric tests on each soil variable independently, testing for an effect of the geographical zone or the rice growing system, using the *kruskal_test* function from the *rstatix* package. Dunn tests (*dunn_test* function) were then performed in case of significant effect evidenced to identify statistically differing groups.

For microbial communities, PERMANOVA were used to test for significant effects of root compartments, geographical zones and rice growing systems on microbial β-diversity, based on a Bray–Curtis dissimilarity matrix. Sites differences were further tested using the *pairwise.adonis* function of the same *vegan* package. The graphical representation of β-diversity was based on Non-metric Multi-Dimensional Scaling (NMDS, *metaMDS* function). The effect of edaphic variables (i.e., pH, total organic carbon, total phosphorus, total nitrogen, total potassium, CEC and SBE) in structuring β-diversity was tested using the *envfit* function (9999 permutations) in R package *vegan*. Structuring soil properties were thus fitted onto the ordination space.

We tested for an effect of the rice growing system on obtained indices of alpha diversity (Shannon diversity index and observed richness). To this purpose, we performed non-parametric statistical tests (*kruskal_test* function from the library *rstatix*) independently for each kingdom (prokaryotes and fungi respectively) and each compartment (rhizosphere and root associated). In addition, for 16S data only (because of insufficient sample size for ITS), we also tested for an effect of the specific site, using Kruskal-Wallis test, and then performed posthoc tests using *dunn_test* function.

Then, we identified particular soil taxa that were associated with lowlands or irrigated systems using indicator analyses with the function *multipatt* implemented in the *indicspecies* package (De Cáceres *et al*. 2010). The algorithm determines both fidelity and consistency to a system.

Finally, we analyzed the repartition of ASVs assigned to the class *Glomeromycetes* (arbuscular mycorrhizal fungi, AMFs). The summed abundance of all ASVs which best blast hit is within the *Glomeromycetes* class was modeled by generalized linear mixed models (GLMMs), with Poisson distribution, with the package *lme4* (Bates et al. 2015), followed by Type III ANOVA with the package *car* (Fox & Weisberg, 2019). The compartment and the rice growing system were included as fixed effects and the geographical zone as a random effect. In addition, differential repartition analysis was performed using DESeq2 (Love *et al*. 2014) to identify *Glomeromycetes* ASVs showing significant enrichment in one of the two rice growing systems.

R code used to perform the analyses and generate the figures are available upon request.

## Results

### Structure of rice soil properties and rice root associated microbial communities

PERMANOVA analysis performed on the Bray-Curtis distance matrix of soil characteristics to describe the overall soil properties, highlighted no significant influence of the rice growing system, but a differentiation according to the geographical zone, for both physical (F = 6.420, r^2^= 0.448, *p* = 0.006, Fig. 1b) and chemical (F = 4.121, r^2^= 0.346, *p* = 0.026) soil parameters. More precisely, we found no effect of the rice growing system but a significant effect of the geographical zone on the clay and sand contents, as well as total Phosphorus, total Potassium, SBE and CEC (Table S1). Posthoc tests revealed that the geographic zones that differed statistically were the same for the six above-mentioned variables, namely Banzon and Karfiguela zones (Fig. 1c).

To determine whether the geographical zones (Karfiguela, Bama or Banzon), the rice-growing systems (irrigated vs. rainfed lowlands), or their interactions, structured root-associated or rhizosphere microbial communities, we performed PERMANOVA analysis on the Bray-Curtis distance matrix of 16S and ITS ASVs, respectively (Table S2, Fig. S2). As different structures were revealed between root and rhizosphere communities of both prokaryotes (F= 6.863, r^2^ = 0.052, *p* < 0.001) and fungi (F= 3.753, r^2^ = 0.080, *p* < 0.001), we further subsetted both datasets to observe the relative influence of the rice-growing systems and the geographical zones in shaping root and rhizosphere communities separately (Table 1, Fig. 2).

**Figure 2.**
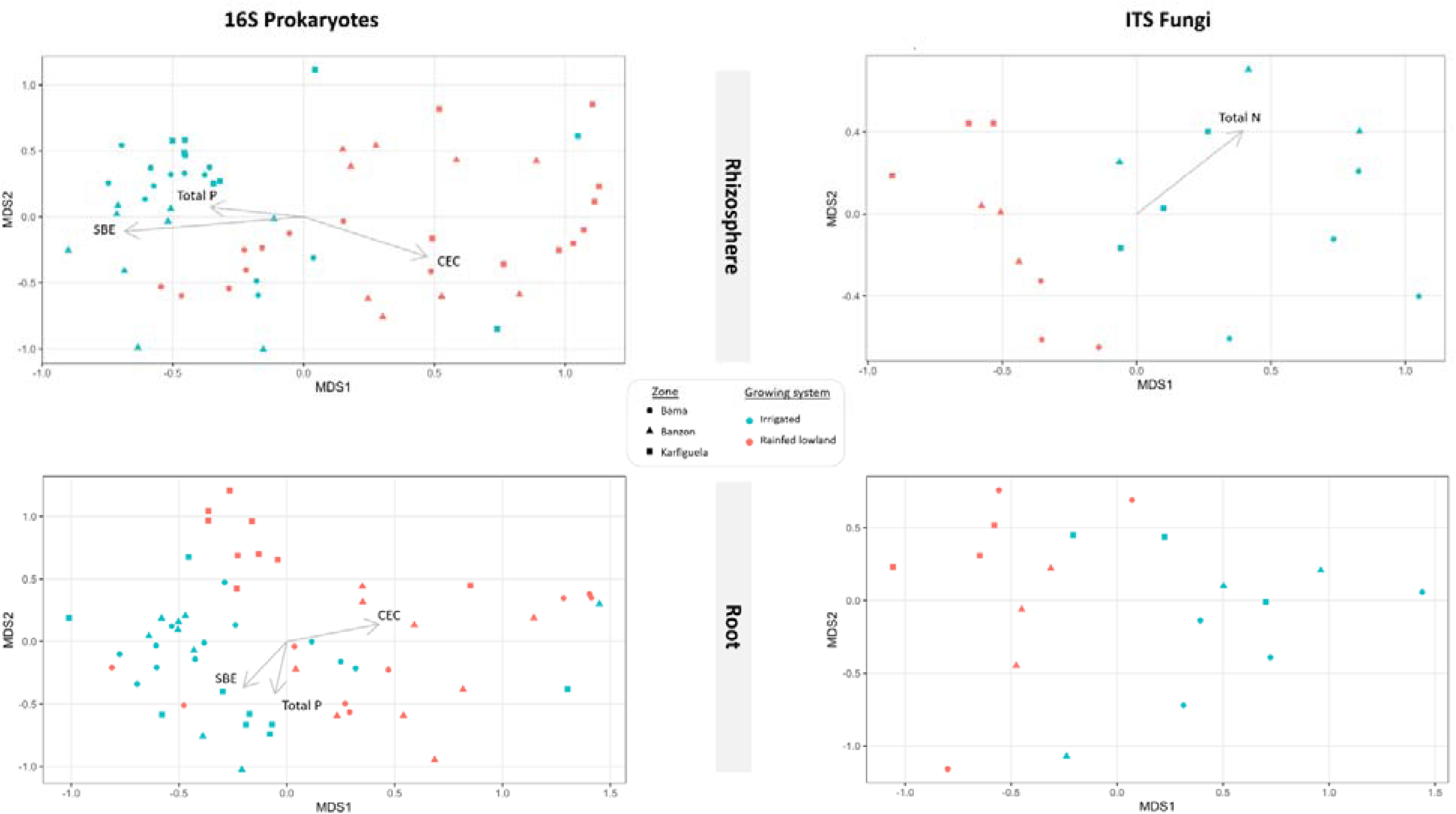
NMDS ordination showing the three factors identified as drivers of the structuration of rice root microbial communities: the color of points represent the rice growing system (irrigated vs rainfed lowland), while the shape shows the geographical zone (Banzon, Karfiguela and Bama). On the left side are presented the analyses based on 16S rRNA gene reflecting Prokaryote communities, where one point corresponds to one plant. On the right side are shown the analyses based on ITS reflecting fungal communities, where one point corresponds to one field. The root compartment is presented on the upper side of the figure while the rhizosphere data appear on the bootom side. Only the soil physicochemical parameter that revealed as having a significant effect (see Supplementary Table S4) are represented with arrows: cation exchange capacity (CEC), Sorptive Bioaccessibility Extraction (SBE), total concentrations of phosphorus (Total P), total concentrations of nitrogen (Total N), total organic carbon (Organic C).

**Table 1.**
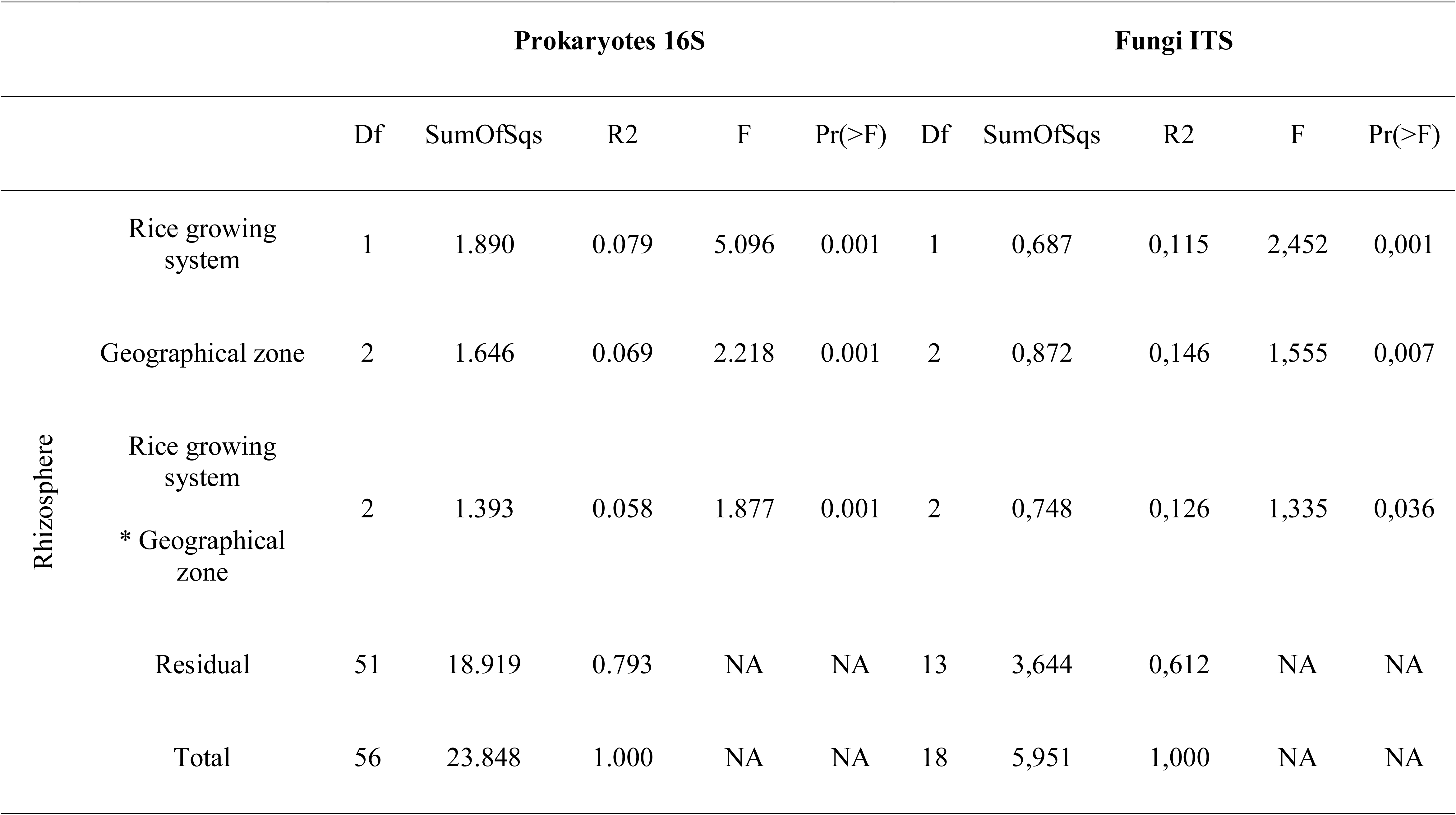
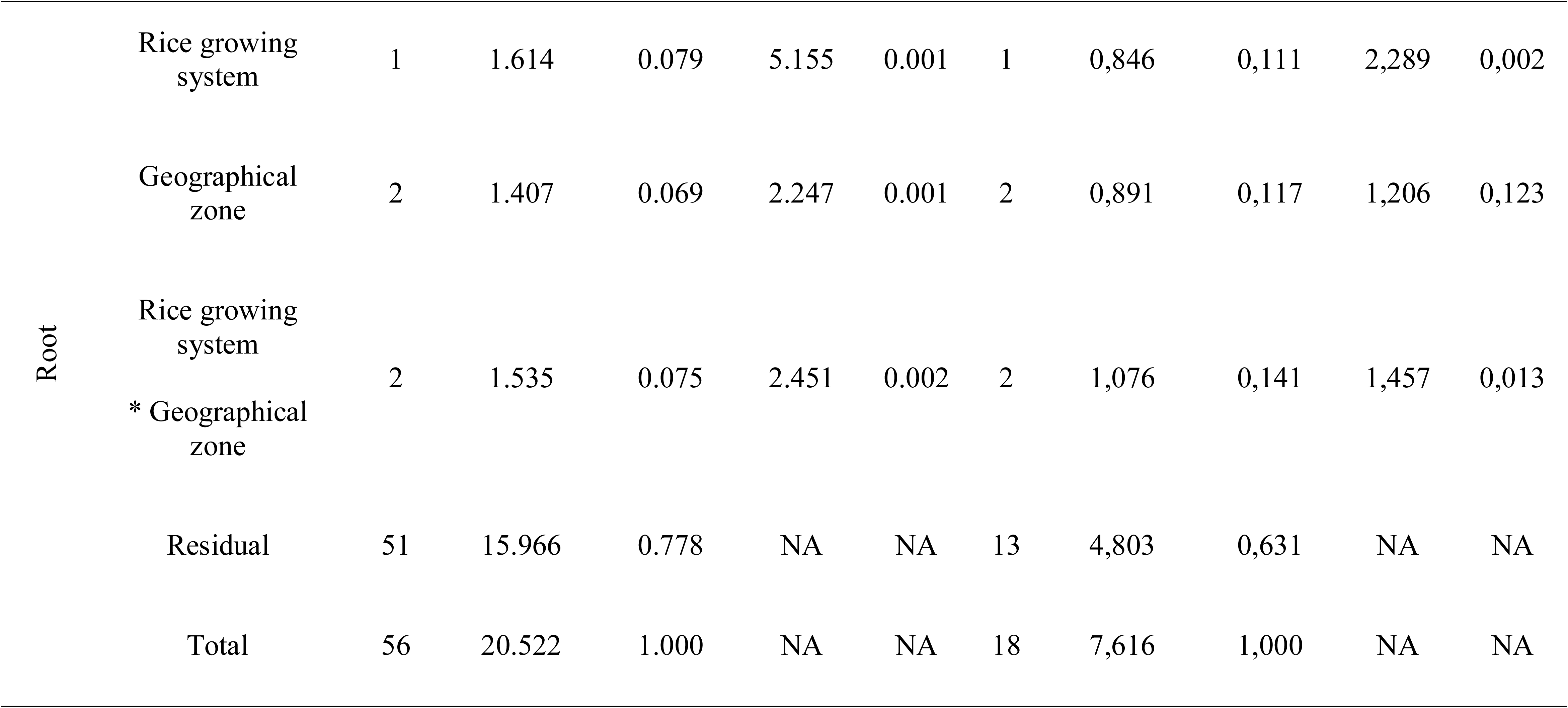
Results of PERMANOVA analysis performed independently for rhizosphere compartment and root compartment, for 16S and ITS microbiome data.

In the rhizosphere, prokaryotic communities were mainly structured by the rice growing system (F=5.096, r^2^ = 0.079, *p* < 0.001, Table 1), the geographical zone (F=2.118, r^2^ = 0.069, *p* < 0.001) and the interaction between rice growing system and zone (F=1.877, r^2^ = 0.058, *p* < 0.001). Posthoc tests revealed that most of the pairs of sites (i.e., irrigated and lowland from the same geographical zone) were significantly different, but interestingly revealed no significant difference between communities originating from irrigated systems, whereas all communities from rainfed lowland sites exhibited distinct structures (Table S3). Fungal communities of the rhizosphere were also mostly structured by the rice growing system (F=2.452, r^2^ = 0.115, *p* < 0.001, Table 1). The geographical zone (F=1.555, r^2^ =0.146, *p* = 0.007) and the interaction between rice growing system and the geographical zone (F=1.335, r^2^ = 0.126, *p* = 0.036) were also driving the rhizosphere fungal microbiome. The low number of samples did not allow to detect, if any, statistically significant differences between sites in communities’ structures for ITS (Table S3).

As observed for rhizosphere communities, the root-associated prokaryotic communities were mainly shaped by the rice growing system (F=5.155, r^2^ = 0.079, *p* < 0.001, Table 1), the interaction between rice growing system and the geographical zone (F=2.451, r^2^ = 0.075, *p* = 0.002) and the geographical zone (F=2.247, r^2^ = 0.069, *p* < 0.001). Most of the pairs of sites (from the same geographical zone) were significantly different. No significant difference was detected between communities originating from irrigated systems, whereas all communities from rainfed lowland sites exhibited distinct structures (Table S3). Root-associated fungal communities were also mostly influenced by the rice growing system (F=2.289, r^2^ = 0.111, *p* = 0.002, Table 1), and by the interaction between rice growing system and the geographical zone (F=1.457, r^2^ =0.141, *p* = 0.013). The effect of the geographical zone on root-associated fungal communities was not evidenced (F=1.206, r^2^ = 0.117, *p* =0.123). As for rhizosphere, posthoc tests on root-associated fungal communities were all non-significant (Table S3).

The influence of soil chemical parameters on microbial community structure is reported in Fig. 2 as arrows and in Table S4. We noticed that the prokaryotic communities of both rhizosphere and roots were affected by the same three parameters: SBE (r^2^ =0,482, p < 0,001 for rhizosphere, and r^2^ = 0,175, p = 0,006 for roots), CEC (r^2^ =0,314, p < 0,001 for rhizosphere, and r^2^ = 0,204, p = 0,003 for roots) and total phosphorus (r^2^ = 0,132, p = 0,023 for rhizosphere, and r^2^ = 0,179, p = 0,004 for roots). For fungi, although various parameters were marginally significant in each compartment (Table S4), we only detected a significant effect of total nitrogen on rhizosphere communities (r^2^ =0,320, p = 0,043).

### Composition of rice root microbiomes and comparison of alpha-diversity

While 16S data were assigned at the genus level for 64% of ASVs, only 34% of ITS ASVs could be assigned (see assignations at the phyla level in Fig. S4). Assignations at the phylum level were obtained for all (100%) 16S ASVs, but only for 62% for ITS ASVs (see Fig. S4). Assigned prokaryotic taxa represent 17 phyla, most abundant ones being Proteobateria, Firmicutes, Mixoccoccota and Acidobacteriota. For ITS, seven phyla were found, with the most abundant ones being Ascomycota followed by Basicomycota.

We tested the effect of the rice growing system on the diversity indices (alpha-diversity). No effect could be evidenced on the fungal (Shannon: H = 0.026, p =0.87 for the rhizosphere, and H = 0.107, p =0.74 for root-associated communities), or root-associated prokaryote diversities (Shannon: H = 1.05 p = 0.306). On the other hand, the rice growing system had a significant effect on the prokaryote diversity of the rhizosphere (Shannon: H = 11.6, p <0.001), with a higher Shannon diversity index in irrigated areas (5.03 ± 0.13), compared to rainfed lowlands (4.39 ± 0.12) and a higher observed richness (275.6 ± 26.4) in irrigated areas compared to rainfed lowlands (143.7 ± 17.4) (Fig. 3). We noticed however an opposite pattern for fungal communities of the rhizosphere with higher observed richness in rainfed lowlands, compared to irrigated areas (Fig. 3).

**Figure 3.**
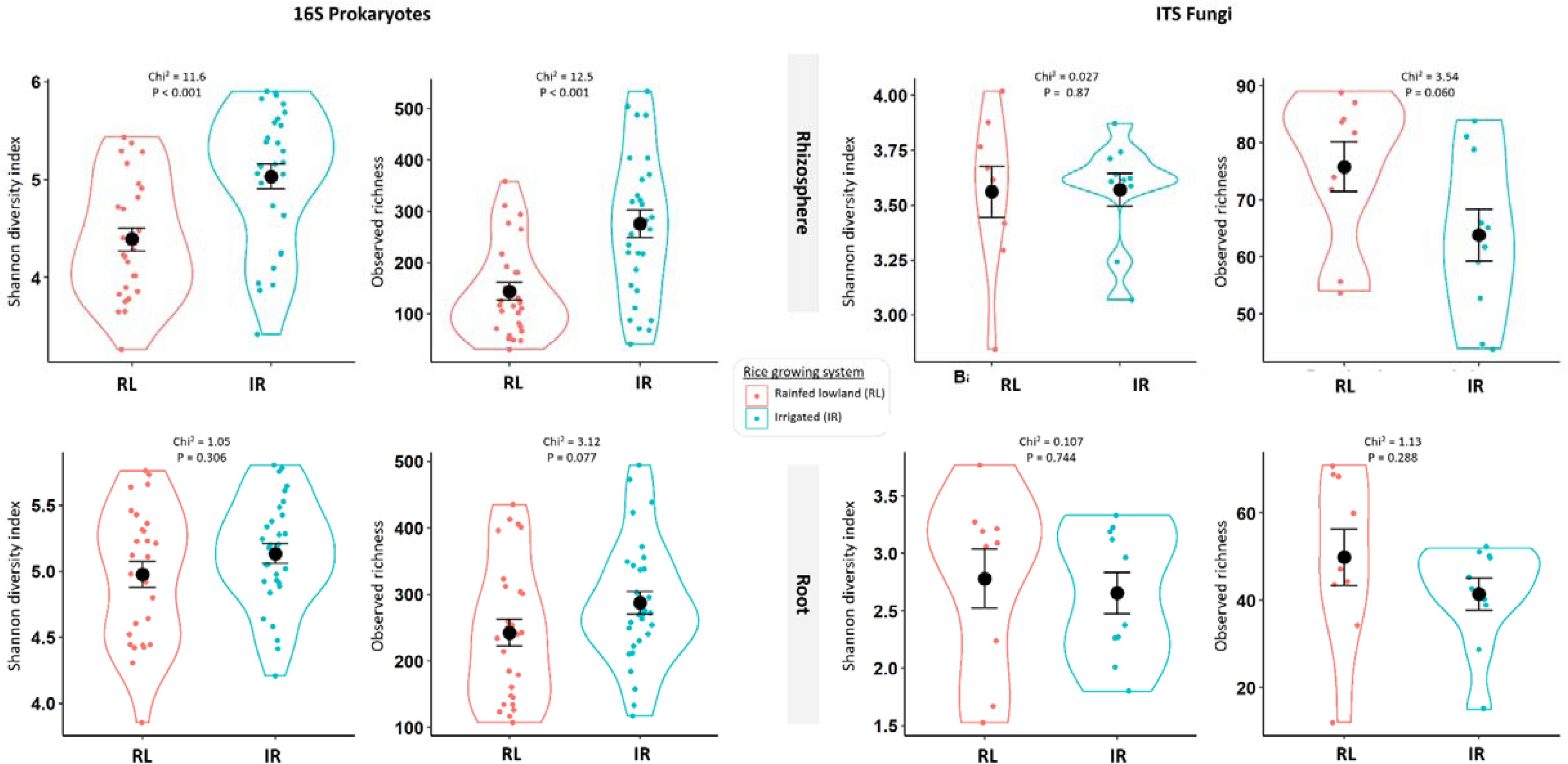
Comparison of root-associated microbiota α-diversity in contrasted rice-growing systems: irrigated (in blue) vs rainfed lowland (in red). Observed richness and Shannon indices are reported for each sample (i.e. one plant for 16S and one field for ITS), as violin diagram for each rice growing system. The left side of the figure presents the results obtained for 16S microbiome, while the right side shows the results obtained for ITS analysis. On top are shown the results of the rhizosphere compartment and on the bottom are the results obtained for the root associated compartment.

This effect of the rice growing system on 16S rhizosphere data was also clearly observed when plotting the diversity indices by site (Fig. S5). In addition, we found that the specific site had an effect on the prokaryotic communities of the rhizosphere, and also, but to a lesser extent, in roots (Fig. S5 and Table S5). In the rhizosphere, the highest diversity was found in the irrigated perimeter of Karfiguela, and to a lesser extent in the irrigated area of Bama. A particularly low diversity was found in the rainfed lowland of Karfiguela zone, and to a lesser extent in the rainfed lowland of Banzon. Conversely, we noticed a slightly higher diversity in fungal root associated communities in the rainfed lowland of Karfiguela (Fig. S5).

### Core microbiome and co-occurrence networks in the two rice-growing systems

ASVs belonging to the core microbiome of lowland vs. irrigated rice were respectively identified with a prevalence threshold set to 60%. For 16S, we identified 26 core ASVs associated with the irrigated systems, and two core ASVs in lowlands (Fig. 4). Among the core taxa in irrigated areas, the vast majority of phylotypes (25/26) belonged to the *Burkholderiaceae* family, with 24 assigned to *Ralstonia pickettii* and one to *Paraburkholderia kururiensis*. One of the core ASVs is common to both irrigated area and rainfed lowlands systems. Its best blast hit corresponds to *Bradyrhizobium tropiciagri* (Bradyrhizobiaceae) with a 99.5% sequence similarity. Another core phylotype in rainfed lowlands is assigned to the same species with 99.3% sequence similarity.

**Figure 4.**
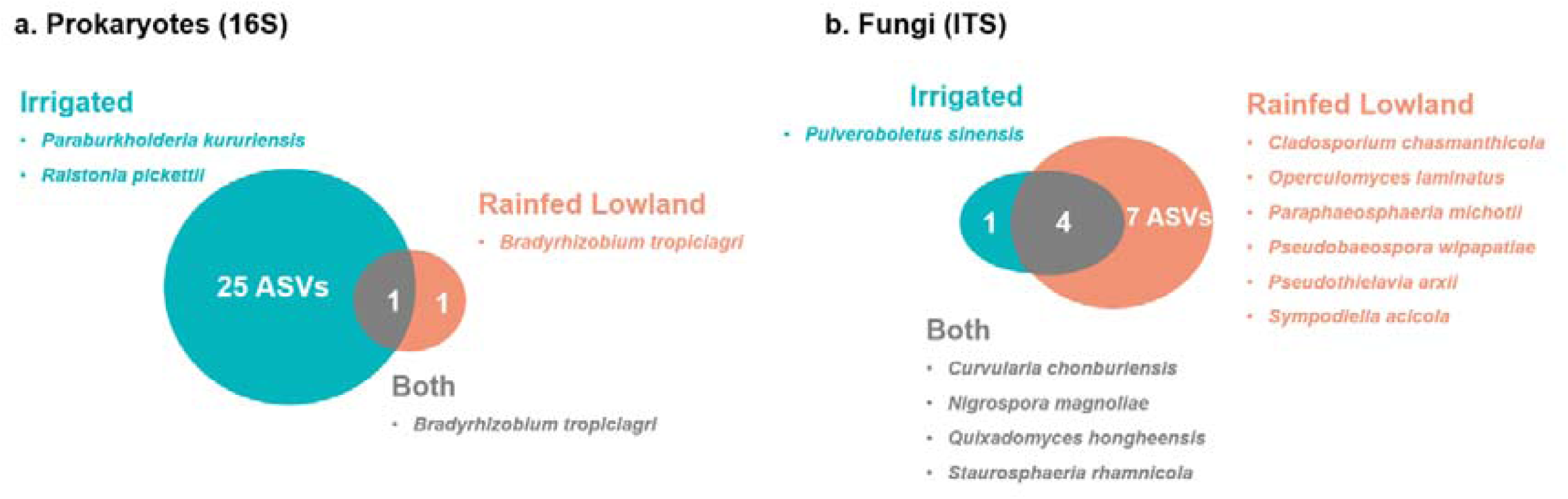
Venn diagram representing the core sequence variants for each rice growing system : irrigated vs rainfed lowlands. A. For prokaryotes B. For fungi

For ITS, we identified 5 core ASVs in the irrigated systems, compared to 11 core ASVs associated with the lowlands, 4 of them being common to both rice growing systems (Fig. 4).

Then, we compared the prokaryotic co-occurrence networks in each rice growing system respectively (Table 2). We identified 15 hub ASVs in the irrigated systems and 20 in rainfed lowlands. We found a higher edge number in irrigated compared to rainfed lowlands: 1720 positive and 269 negative resulting in 2029 total edges in irrigated areas, while only 1163 positive and 85 negative resulting in 1248 total edges were found in rainfed lowlands. Finally, the network computed from irrigated areas had higher connectivity compared to the one from rainfed lowlands (9.8 vs 7.9 node mean degrees, respectively).

**Table 2.**
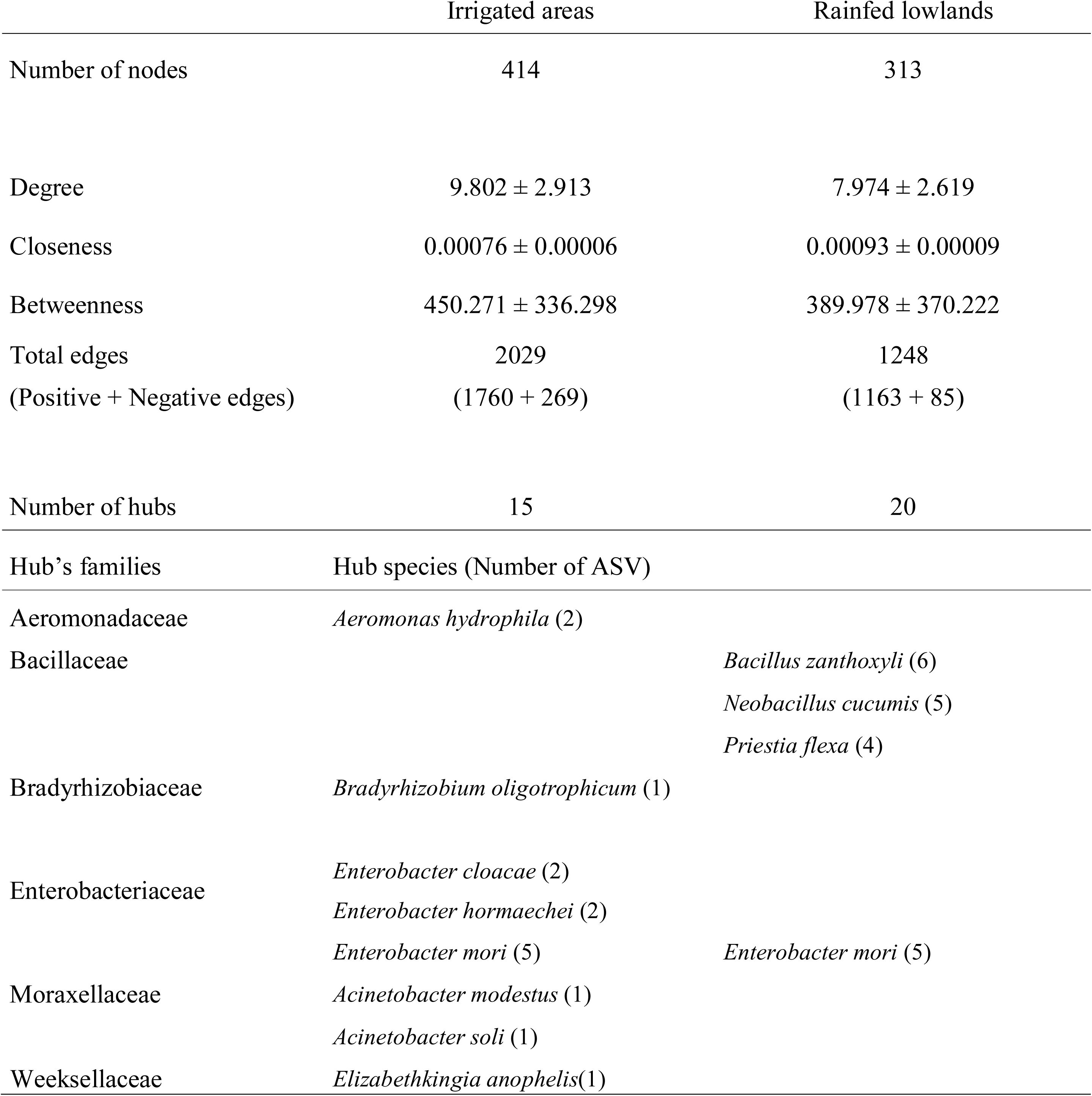
Properties of co-occurrence networks of Prokaryote taxa in rhizosphere and root-associated samples from irrigated areas in the one hand, and rainfed lowlands in the other hand.

None of the identified hub taxa were also core in any of the two rice growing systems. Only one ASVs was identified as a hub in both irrigated and rainfed lowland systems, assigned to *Enterobacter mori* (Enterobacteriaceae). Hub taxa in irrigated areas (15 ASVs) were assigned to 8 different species from 5 families, while hub taxa in rainfed lowland (20 ASVs) only corresponded to 4 species from 2 families (Table 2).

### Indicator taxa of the two rice growing systems

For 16S data, we found 128 indicator taxa in irrigated areas, including ASVs from eight bacterial families, most of them assigned to *Acinetobacter, Ralstonia, Aeromonas, Comamonas, Clostridium* and *Enterobacter* (Table 3). On the other hand, only 63 were identified in rainfed lowlands, most of them within the Bacillaceae family, including ASVs assigned to *Exiguobacterium* and *Priestia*, and Bradyrhizobiaceae family, genus *Bradyrhizobium* (Table 3). The ASV assigned to *Paraburkholderia kururiensis* (Burkholderiaceae) revealed as indicator in irrigated areas (Table 3) was also a core taxa in irrigated areas. Also, among the 24 indicator ASVs in irrigated areas assigned to *Ralstonia pickettii* (Burkholderiaceae), 16 were also core in irrigated areas. In addition, four indicator ASVs in irrigated areas were also hubs in this system: two assigned to *Aeromonas hydrophilai* (Aeromonadaceae), another assigned to *Enterobacter cloacae* (Enterobacteriaceae), and finally one corresponding to *Acinetobacter soli* (Moraxellaceae). Three ASVs assigned to *Priestia flexa* (Bacillaceae) were hubs in rainfed lowlands.

**Table 3.**
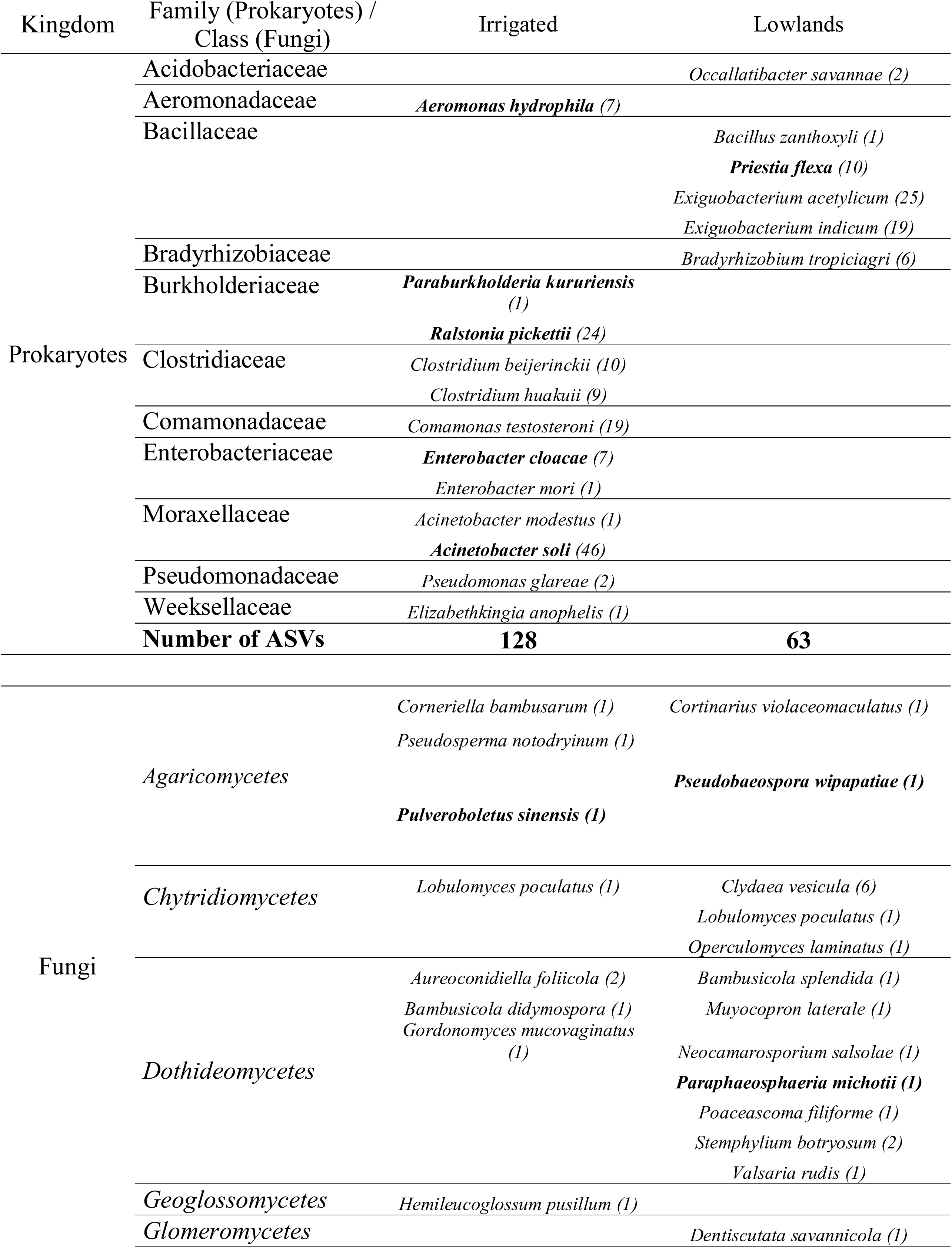
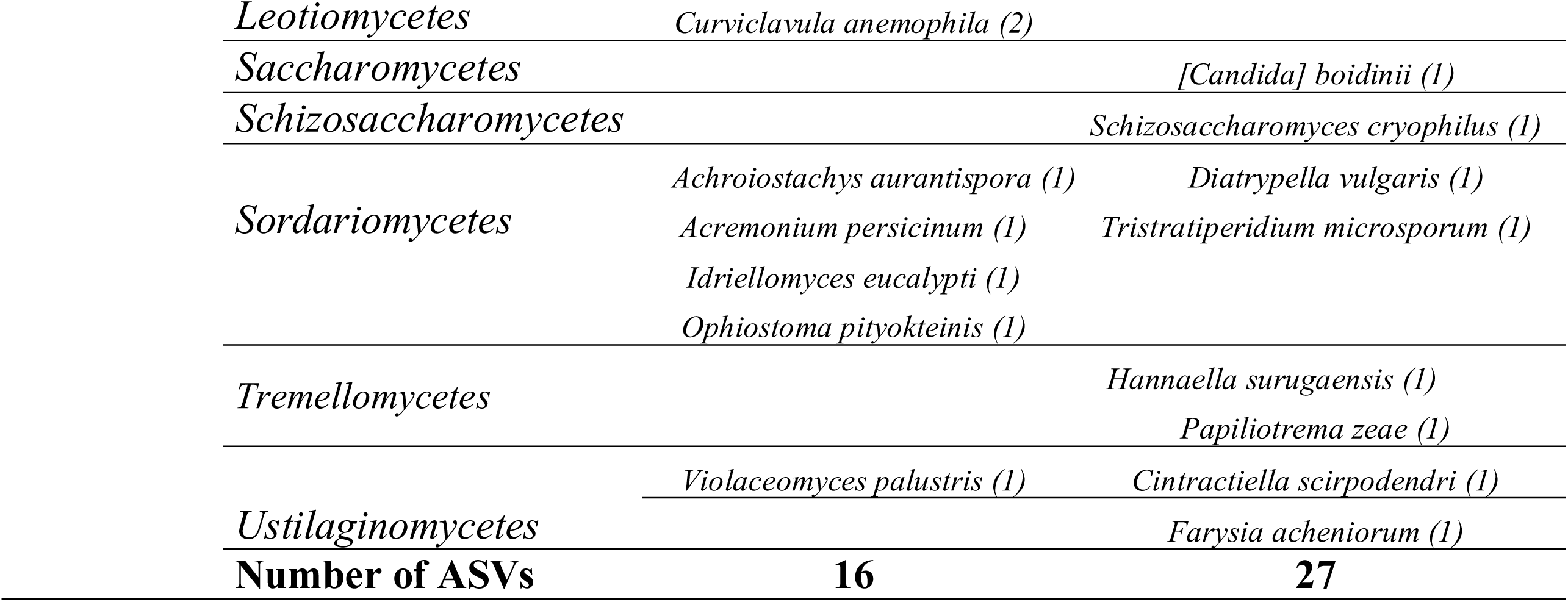
List of species assignation and number of sequence variants (ASVs) identified as indicator taxa in irrigated and rainfed lowland environment. The species in bold were also found as potential hub or core taxa.

For ITS data, we found 16 indicator taxa in irrigated areas, and 27 in rainfed lowlands (Table 3). Indicator taxa in irrigated areas were assigned to seven classes: *Agaricomycetes*, *Chytridiomycetes*, *Dothideomycetes*, *Geoglossomycetes*, *Leotiomycetes*, *Sordariomycetes*, and *Ustilaginomycetes*. Indicator taxa in rainfed lowlands were assigned to nine classes: *Agaricomycete*s, *Chytridiomycetes*, *Dothideomycetes*, *Glomeromycetes*, *Saccharomycetes Schizosaccharomycetes*, *Sordariomycetes*, *Tremellomycetes*, *Ustilaginomycetes*. One ITS ASV identified as indicator taxa in irrigated, with best hit *Pulveroboletus sinensis* (Agaricomycetes), was also core in this rice growing system, and two indicator taxa in rainfed lowlands were also core in this system: one assigned to *Pseudobaeospora wipapatiae (*Agaricomycetes) and the other to *Paraphaeosphaeria michotii* (Dothideomycetes).

### Putative pathogen or phytobeneficial taxa

First, responses for an ‘*oryz*’ query within assignation and blast, found matching records only in the 16S dataset. A total of 200 ASVs included ‘*oryz’* in their names, from 21 different genera, none of these species corresponded to pathogens from Table S6. Among them, putative beneficial taxa were found, particularly the following: *Azospirillum oryzae, Novosphingobium oryzae, Paenibacillus oryzae,* and *Rhizobium oryzae, R. rhizoryzae and R. straminoryzae*.

Next, we made a subset of the ITS dataset for ASVs assigned to the *Glomeromycetes* class (total of 14 ASVs). AMF summed abundance was affected by the compartment (χ^2^= 101.22, *p*<0.001) and by the rice growing system (χ^2^ = 951.12, *p*<0.001), with higher abundances in rhizosphere compartment and in rainfed lowlands (Fig. 5). In addition, differential abundance testing between rice growing systems detected an ASV assigned to *Racocetra crispa* as preferentially found in rainfed lowlands (l2FC = 24.59; p<0.001). We also noticed that another *Glomeromycetes* (*Dentiscutata savannicola*) was identified as indicator taxa in rainfed lowland environments (Table 3).

**Figure 5.**
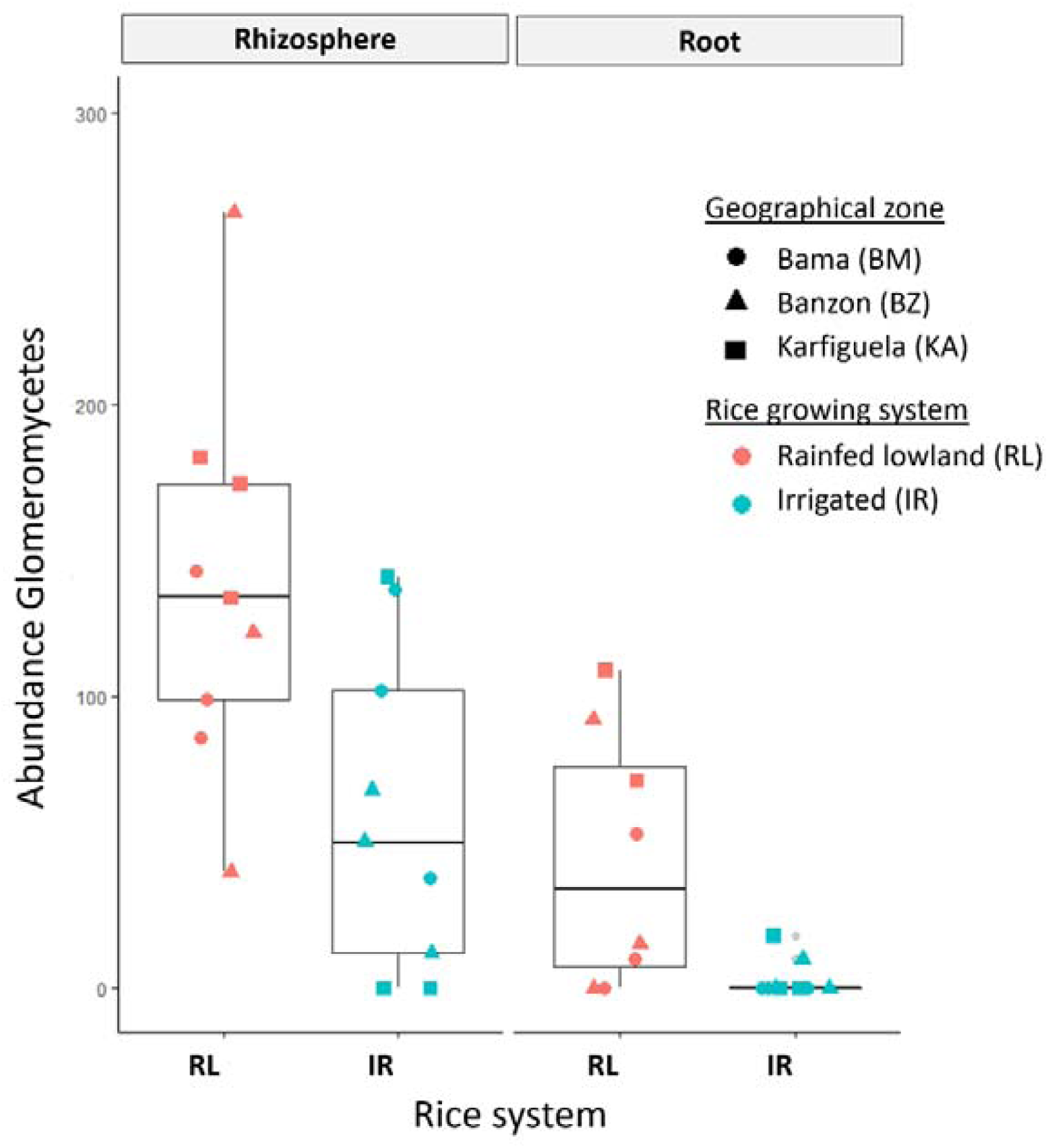
Spatial repartition of summed abundance of 14 ITS ASVs assigned to the class *Glomeromycetes* in the two rice root microbiome compartments (rhizosphere and roots), and in each rice growing system (irrigated perimeters vs rainfed lowlands). The shape of the point corresponds to each geographic zone.

We then screened the list of all assigned ASVs for a set of pathogen species defined *a priori* (see the list in the Table S6). For Prokaryotes (16S data), a number of ASVs corresponded to the genera of pathogens, but only *Burkholderia glumae* (two ASVs), *Acidovorax avenae* (four ASVs) and *Dickeya chrysanthemi* (six ASVs) were identified at the species level. These 12 ASVs identified at the species level were however only found in one sample. The *Xanthomonas* genus was found, but with no assignation to *X. oryzae* (instead, assignation to *X. theicola* which is phylogenetically closed to the rice associated *X. sontii* (Bansal *et al*. 2020). A similar situation was observed for the genera *Pseudomonas, Pantoea*, and *Sphingomonas*. The same analysis of putative pathogens for ITS revealed the presence of the following ten genera: *Alternaria*, *Bipolaris*, *Ceratobasidium*, *Curvularia*, *Fusarium*, *Helminthosporium*, *Microdochium*, *Rhizoctonia*, *Sarocladium*. We notice that one ASV whose best blast hit was *Curvularia chonburiensis* was core in both irrigated and rainfed lowlands (Fig. 4).

## Discussion

This study aimed at describing the rice root-associated microbiome by comparing contrasted rice growing systems in farmer’s fields in Burkina Faso. We found that the rice growing system was a structuring factor for rice root-associated microbiomes, and that the diversity of prokaryotic community from the rhizosphere was higher in irrigated areas compared to rainfed lowland. In addition, we identified a number of phylotypes with potential key roles (hub, core, indicators) in the two contrasted systems, as well as putative phytobeneficial and pathogen species. Although the results on fungi (ITS region) must be taken with caution due to a smaller sample size and the poor representation of obtained sequences in available databases, this study shed light on some drivers of assemblage of rice root associated microbial communities in a sparsely documented African system.

### The structuring of microbial diversity is affected by the rice growing system

Although Edwards *et al*. (2018) showed that the root-associated microbiome of distant field sites converge in similarity during the growing season, our study performed at the maturity stage of rice still evidenced some drivers of rice root-associated microbial communities structure. First, as for most rice microbiome studies, we found an effect of the compartment / micro-habitat (Edwards *et al*. 2015; Santos-Medellín *et al*. 2017; Guo *et al*. 2021; Kawasaki *et al*. 2021), and the geographical zone (Edwards *et al*. 2015; Kanasugi *et al*. 2020) on the beta-diversity of rice root-associated microbiome. In addition, our study shows that the contrasted rice-growing systems, namely irrigated perimeters *vs* rainfed lowlands, harbor contrasted rice root-associated microbial communities, both for prokaryotic and fungal communities, and for rhizosphere and root compartment. We notice that the soil physicochemical properties weakly differ between irrigated areas and rainfed lowland, the soil composition was instead mostly affected by the geographical zones. Consequently, we evidence a structuring effect of the rice-growing system that was only slightly related to contrasted soil physicochemical properties. Our results are in line with a previous study comparing microbiomes from two contrasted water management conditions (upland *vs* lowland rice) in controlled settings (a field experiment in northern Italy), which showed differentiation in microbial communities, particularly for root microbiome, and to a lesser extent in soil samples (Chialva *et al*. 2020).

Only a few other studies compared variable water management agricultural systems. Cui *et al*. (2019) showed that irrigation water quality affected bacterial community alpha and beta diversity in maize, with pH and available phosphorus being the major factors shaping microbiome soil composition. Mavrodi *et al*. (2018) evidenced a slight effect of the three seasons of irrigation on the overall diversity within the rhizosphere microbiome in wheat, but significant differences in the relative abundances of specific taxa.

In our study, some of the soil physicochemical parameters affected rice root-associated microbial communities. In particular, CEC and SBE, that reflect soil exchange capacity and bioaccessibility, were the most important soil parameters for the structure of both rhizosphere and root prokaryotic communities. These parameters are not commonly measured, nor identified as important, in other studies of the root-associated microbiome. Our results argue for including them in soil chemical characterization, to investigate whether their impact in microbiome structure is general or not. In addition, phosphorus content significantly structured the prokaryotic communities, both in rhizosphere and roots. Such an effect of phosphorus is known for the rice root associated microbiome (Long & Yao, 2020). On the other hand, the soil chemical parameter evidenced in this study to structure fungal rhizosphere communities was the total nitrogen (N). This is in accordance with a study by Chen *et al*. (2019) showing that nitrogen input drives changes in the microbial root-associated community structure in wheat. Moreover, Wang & Huang (2021) showed the effect of optimized N application on fungal community structure from paddy soils. Kanasugi *et al*. (2020) also evidenced an effect of soil nitrate on rice root fungal communities in Ghana.

### Effect of rice growing system on alpha-diversity and network topology

We found a higher taxonomic diversity in irrigated areas, compared to rainfed lowlands, for prokaryotic communities of the rhizosphere. Our findings differ from Chialva *et al*. (2020)’s results, where the 16S diversity was similar in lowland and upland rice. Chialva *et al*. (2020) also show significantly higher ITS diversity in lowland rice compared to upland, which may relate to the tendency (not significant, maybe as a consequence of the low sample size for ITS) observed here in the rhizosphere.

It is important to note that our sampling was performed in farmer’s fields, while most results, including those of Chialva *et al*. (2020), were obtained in field trials, potentially explaining the differences. Indeed, in our study, various factors, such as rice cultivars, fertilization regime and rotation, exhibit large variability. However, rice genetic diversity was shown to be comparable in irrigated areas and rainfed lowlands (Barro *et al*. 2021), therefore the effect of rice growing system on alpha-diversity of microbiomes could not be attributed to result from difference in terms of genetic diversity of the host plant.

Considering the irrigated areas as systems with more intensive agricultural practices, compared to rainfed lowland, we expected an opposite pattern of microbiome diversity. Indeed, agricultural intensification was shown to reduce microbial network complexity and the abundance of keystone taxa in roots (Banerjee *et al*. 2019). In addition, the fertilization regime is known to have strong impact on root-associated microbiota (Ding *et al*. 2019; Xiong *et al*. 2021). Various studies showed that organic fertilization enhances microbial diversity (Liu *et al*. 2020). For example, recommended fertilization preserved belowground microbial populations, compared to the fertilization mostly used (‘conventional fertilization’) that depressed bacterial diversity, in experiments performed in China (Ullah *et al*. 2020). We considered the irrigated areas as more intensified systems, compared to rainfed lowland, particularly because only irrigated areas allow growing rice twice a year, and because only rainfed lowland sites presented fields with no mineral fertilization at all (Barro *et al*. 2021). We noticed however that organic fertilization remained rare, and its frequency was not drastically affected by the rice growing system. Finally, transplantation was always performed in irrigated areas, while direct sowing was the most common practice in rainfed lowlands.

On the other hand, paddy soils studied in western Burkina Faso (all over the six sites) are particularly poor if compared for example to a study of more than 8 000 soils in Hunan Province (Duan *et al*. 2020), where average organic carbon was 1.972%, compared to 0.922% in our study, total nitrogen was 0.191%, higher than 0.072%, and total phosphorus was 0.71g.kg^-1^, compared to 0.24g.kg^-1^. The studies previously cited evidencing fertilization effects, were performed in soils with higher carbon and nitrogen contents (see for example Ullah *et al*. 2020, where minimum average organic carbon was 2% and total nitrogen 0.1%). The effect of fertilization on microbial diversity may actually depend on various aspects, including the soil type. Notably, a positive relationship was found between rice fertilization and soil bacterial richness and diversity in a 19-years inorganic fertilization assay in a reddish paddy soil in southern China (Huang *et al*. 2019); while Wang & Huang (2021) showed an effect of the fertilization on paddy soils microbial community composition but no effect on the diversity. In poor soil systems such as in this study, fertilization input may actually increase microbial diversity.

Finally, a complementary hypothesis could be the higher fragmentation of rainfed lowlands compared to irrigated areas. Indeed, irrigated sites correspond to larger areas cultivated in rice, possibly with two rice seasons per year, so that rice fields are likely to be more connected to each other than in rainfed lowlands. Higher connectivity generally leads to higher biodiversity (Fletcher *et al*. 2016). The principles of metacommunity theory could also be applied to micro-organisms, with reduction in host habitats and fragmentation potentially increasing extinction rates (Mony *et al*. 2020), but as our study misses an explicit characterization of the rice landscape structure, this hypothesis could not be formally tested.

### Distant rainfed lowlands differ more than distant irrigated perimeters

Our results showed that the prokaryotic communities in the rice rhizosphere and roots from the three irrigated sites do not differ significantly from each other. On the other hand, the same analysis revealed significant differentiation between the three rainfed lowland study sites (in all three cases for rhizophere and two out of three comparisons in roots). Also, we found very few core phylotypes in rainfed lowland, with only two core ASVs for 16S, what reinforces the above-mentioned observation. These results are likely driven by a higher heterogeneity between rainfed sites, in terms of water control, agricultural practices or rice genotypes.

Indeed, in irrigated rice, the farmer has the potential to control irrigation water during the whole growing season. On the other hand, irrigation in rainfed lowland is dependent on precipitations that differ between the three geographical zone sampled within the rice growing season. In addition, we showed a high heterogeneity of agricultural practices in rainfed lowlands: for example, legume rotation was common in the rainfed lowland of Bama zone, but rare or absent in the two other rainfed lowland sites, and organic fertilization was more frequent in the rainfed lowland of Karfiguela zone, than in the other sites (Barro *et al*. 2021). Finally, in terms of rice genetics, a high rice genetic differentiation was found between the rainfed lowland site of Karfiguela zone and the five sites: a distinct genetic group *O. sativa Aus*, and other distinct landraces were found in this peculiar site, compared to the five others where only *O. sativa indica* was grown (Barro *et al*. 2021). These specificities of the rainfed lowland from Karfiguela zone, in terms of rice grown and agricultural practices, may also drive its specific patterns of alpha diversity, with a particularly low prokaryote diversity (in rhizosphere and also, in a lesser extent, in roots), and a tendency for higher fungal diversity in roots.

Our sampling size was much lower for ITS and this likely explains the absence of such a pattern, with no significant differences obtained between pairs of sites. Alternatively, the pattern may be different for fungal diversity, as suggested by the higher number of core taxa in rainfed lowlands than in irrigated areas.

### Identification of core microbiota and hub phylotypes

The prevalent taxa, indicator taxa and hubs may be considered as having an important ecological role in microbiome assembly and ecosystem functions (Banerjee *et al*. 2018). In this study, we identified the core prokaryote and fungal microbiota in both irrigated and rainfed lowland environments. While four fungal taxa were found to be cores in both systems, only one bacterial core taxa was shared between the two rice growing systems: assigned to *Bradyrhizobium tropiciagri*, a nitrogen-fixing symbiont isolated from tropical forage legumes (Delamuta *et al*. 2015), known as rice root endophytes capable of fixing N_2_ (Chaintreuil *et al*. 2000; Ding *et al*. 2019). In addition, the core taxa in irrigated areas likely includes *Paraburkholderia kururiensis*, a bacterium with potential phytobeneficial properties (bioremediation, biofertilization and biocontrol of pathogens; Dias *et al*. 2019). Various ASVs identified as core in irrigated areas were assigned to *Ralstonia pickettii*, an ubiquitous Betaproteobacteria found in water and soil, and capable to thrive in low nutrient (oligotrophic) conditions (Ryan *et al*. 2007). Isolated from the plant rhizosphere, *R. pickettii* injected in tomato stem could reduce bacterial wilt disease caused by its congeneric pathogen *R. solanacearum*, and could consequently be considered as potential biocontrol agent (Wei *et al*. 2013). Noteworthy, this bacterium is also described as human emerging pathogen, causing nosocomial infections (Ryan *et al*. 2006).

One ASVs assigned to *Enterobacter mori* (Enterobacteriaceae) was identified as hub of the 16S-based co-occurrence networks both in irrigated areas and rainfed lowland site. On the other hand, most of the bacterial taxa identified as hubs differed between irrigated areas and rainfed lowlands reflecting a highly contrasted structuring of bacterial communities in the two rice growing systems.

### Characterization of indicator taxa in each rice growing system and contrasted repartition of AMFs

We identified indicator taxa for each rice growing system, most of them being in irrigated for procaryotes (128, *vs* only 63 in rainfed lowlands) while the opposite was found for fungi (27 in rainfed lowlands *vs* only 16 in irrigated areas). For prokaryotes, five taxa identified as indicator species in irrigated areas were also core or hub: the previously mentioned *Paraburkholderia kururiensis* and *Ralstonia pickettii* as well as *Aeromonas hydrophila*, *Enterobacter cloacae* and *Acinetobacter soli*. In rainfed lowlands, it was the case for *Priestia flexa*. In particular, we notice that *Acinetobacter soli* was identified as potent phosphorus solubilizer in rice and consequently promising for plant growth promotion (Rasul *et al*. 2019).

For fungi, we identified as both indicator and core taxa: *Pulveroboletus sinensis* in irrigated areas, as well as both *Pseudobaeospora wipapatiae* and *Paraphaeosphaeria michotii* in rainfed lowlands. Eight ASVs assigned to *Chytridiomycetes* were identified as indicator species in rainfed lowland systems. Members of this fungal division of aquatic fungi (Barr, 2001) were found in the rhizosphere compartment in our study. They are known as particularly abundant in microbial communities associated with rice roots, compared to other crops (Ding *et al*. 2019) and were preferentially associated to lowland conditions compared to upland (Chialva *et al*. 2020).

We identified a few potentially beneficial taxa that could be investigated further. In particular, AMFs of the class *Glomeromycetes* were found preferentially in rainfed lowlands, with one ASV, *Racocetra crispa,* enriched in rainfed lowland system compared to irrigated areas, and one ASV, *Dentiscutata savannicola*, identified as indicator in rainfed system. This was expected considering the lower frequency of mineral fertilization in rainfed lowlands compared to irrigated areas. Indeed, AMF colonization was shown to be affected by farming regimes: the rice roots cultivated in the conventional agrosystem (N and P fertilization and pesticides) or under permanent flooding showed no AMF colonization, while the rice plants grown with organic conditions showed typical mycorrhization patterns (Lumini *et al*. 2011).

### Towards the identification of putative pathogens and the study of interactions between microbiome and diseases

Various pathogen species are suspected from the sequence variants identified in this study. In particular, the 16S dataset contained ASVs assigned to *Burkholderia glumae*, *Acidovorax avenae* and *Dickeya chrysanthemi*. All three remained rare, with only one sample containing each of these sequences. The presence of *B. glumae* was described in Burkina Faso (but with no molecular data; Ouedraogo *et al*. 2004), and targeted detection performed in two sites failed to detect *B. glumae* and *A. avenae* (Bangratz *et al*. 2020). For fungal pathogens, we found ASVs assigned to ten genera comprising rice pathogens, including *Bipolaris*, *Curvularia*, *Fusarium* and *Rhizoctonia*. It has to be noted that the major rice pathogens causing foliar diseases, namely *Pyricularia oryzae* and *Xanthomonas oryzae*, known from symptom observations to be present in the study sites (Barro *et al*. 2021), were not detected in this root-associated metabarcoding data. Some of these putative fungal pathogens are frequent, particularly one, whose best blast hit is *Curvularia chonburiensis*, identified as core taxa in both irrigated perimeters and rainfed lowlands. Various *Curvularia* species are known to be pathogenic in rice, potentially causing contrasted symptoms (Gao *et al*. 2012; Majeed *et al*. 2015), and their widespread repartition evidenced here argues for more work in plant pathology to better understand the interactions between *Curvularia* and rice.

The literature shows that higher microbiome diversity may be associated with a lower infection rate (see for example Rutten *et al*. 2021). Our results for rice from western Burkina Faso are somehow opposite as irrigated perimeters harbor more diverse prokaryotic communities of rhizosphere compared to rainfed lowlands (this study), but also higher prevalence of major rice diseases, particularly bacterial leaf streak and the fungal rice blast disease, based on the observation of foliar symptoms (Barro *et al*. 2021). On the other hand, for fungal communities, the pattern may actually be opposite; a tendency for higher diversity in rainfed lowlands is observed but low sample size prevents from obtaining significant results. Also, some diseases, such as the viral yellow mottle disease (Barro *et al*. 2021), are not affected by the rice growing system but by the specific site. The relationship between the diversity of root-associated microbiome and diseases is complex and remains to be studied in more details, including under controlled conditions.

More generally, scientific interest in the relationship between root-associated microbiota and plant diseases is growing (Vannier *et al*. 2019; Trivedi *et al*. 2020). In rice, various studies evidenced an inhibition of disease development by root-associated micro-organisms (see Yasmin *et al*. 2016 for bacterial diseases; and Spence *et al*. 2014; Law *et al*. 2017 for rice blast). On the other hand, disease was shown to affect root-associated microbiomes, with for example the effect of *Magnaporthe grisea* inoculation on microbial endosphere diversity (Tian *et al*. 2021). A promising avenue of research is consequently to investigate the relationship between root-associated microbiota and rice diseases in the particular rice-growing systems of Burkina Faso.

### Perspectives

We are only at the beginning of understanding the complexity of rice root microbial communities, especially for rice cultivation in Africa. The originality, but also a limitation of our study, lies in the fact that the samples were collected in farmer fields, and it globally compares the two contrasting rice production systems that differ in various management practices, so that it could not tease apart the specific effect of each individual factor (water management, variety, fertilization, pesticides, etc). More investigations are now required to decipher each structuring factor at a smaller scale: in particular between fields within each site, where the rice cultivar and specific agricultural practices are likely to play a significant role (Delitte *et al*. 2021).

Describing rice microbiota through metabarcoding is a first mandatory step that needs to be combined with culturomics for a greater accuracy and a deeper description, in particular in such systems where some taxa are poorly described in taxonomic databases. Experimental work in an integrative approach is also required to move on towards microbiota management methodologies. Such microbiota-based strategies could contribute to improving rice health and productivity (Sessitsch & Mitter, 2015), while preserving human health. They are consequently an important component of the toolbox of science-based strategies to achieve zero-hunger in Africa.

## Acknowledgments

This work was performed thanks to the facilities of the “International joint Laboratory LMI PathoBios: Observatory of plant pathogens in West Africa: biodiversity and biosafety” (www.pathobios.com; twitter.com/PathoBios), and conducted within E-SPACE project (www6.inrae.fr/e-space/), led by Claire Neema. We are very grateful to Abdoul Kader Guigma, Yacouba Kone, Moumouni Traoré and Arzouma Diasso for their contributions to the fieldwork in Burkina Faso. We thank the rice farmers from Badala, Bama, Senzon, Banzon, Tengrela and Karfiguela for their kind collaboration. We thank Corinne Vacher and Alain Sarniguet for constructive discussions at earlier stage of the project. The manuscript benefited from helpful comments by Philippe Roumagnac, who also contributed to the initial design of the study. Finally, we dedicate this manuscript to the late Christian Vernière for his kind feedback at different stages of this project.

---

**Figure S1:**
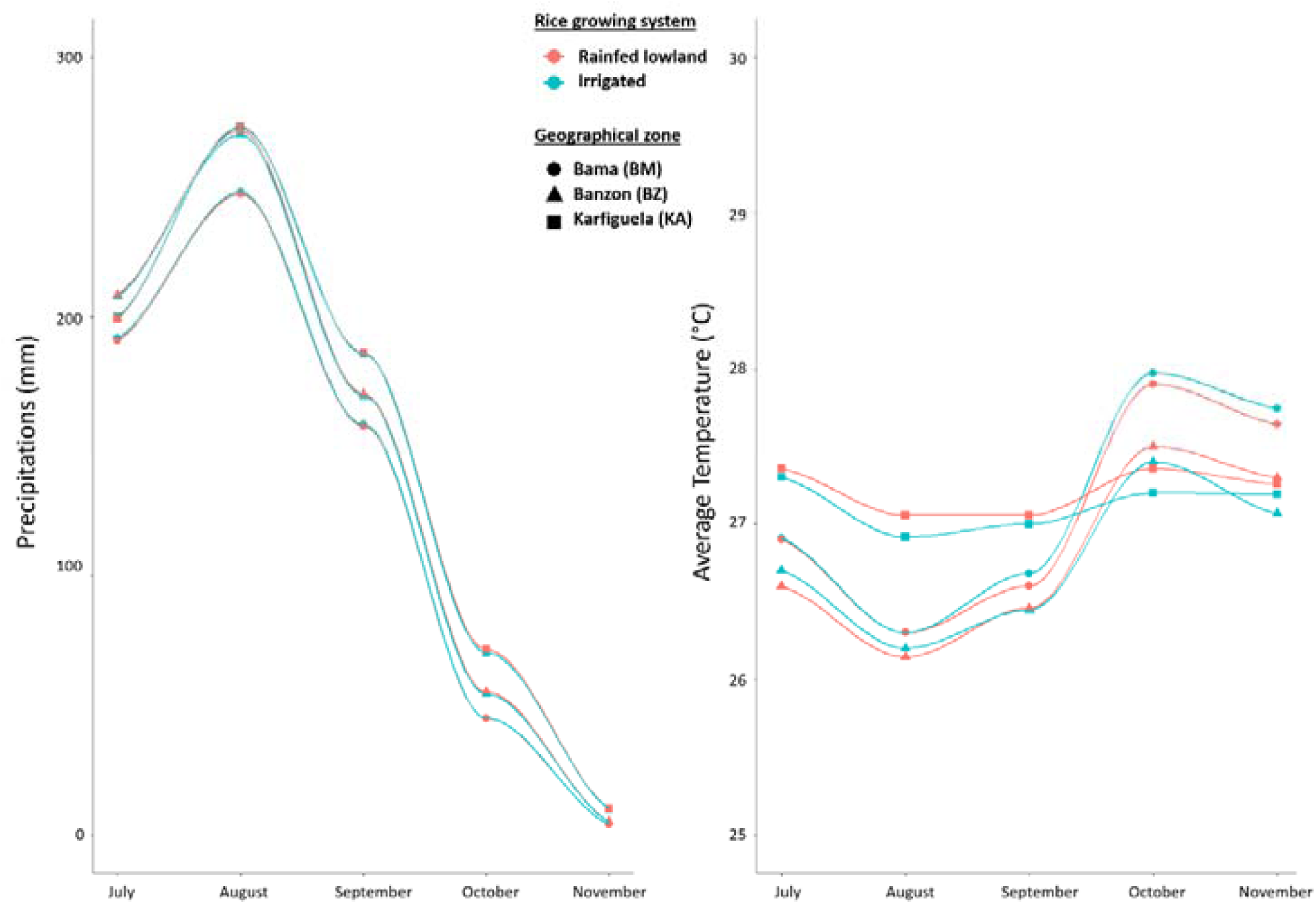
Monthly precipitation (on the left) and average temperature (on the right) over 1970-2000 period, for each of the six study sites in Western Burkina Faso during the rice growing season (July to November). WorldClim 2 data (Fick & Hijmans, 2017): https://worldclim.org/data/worldclim21.html

**Figure S2:**
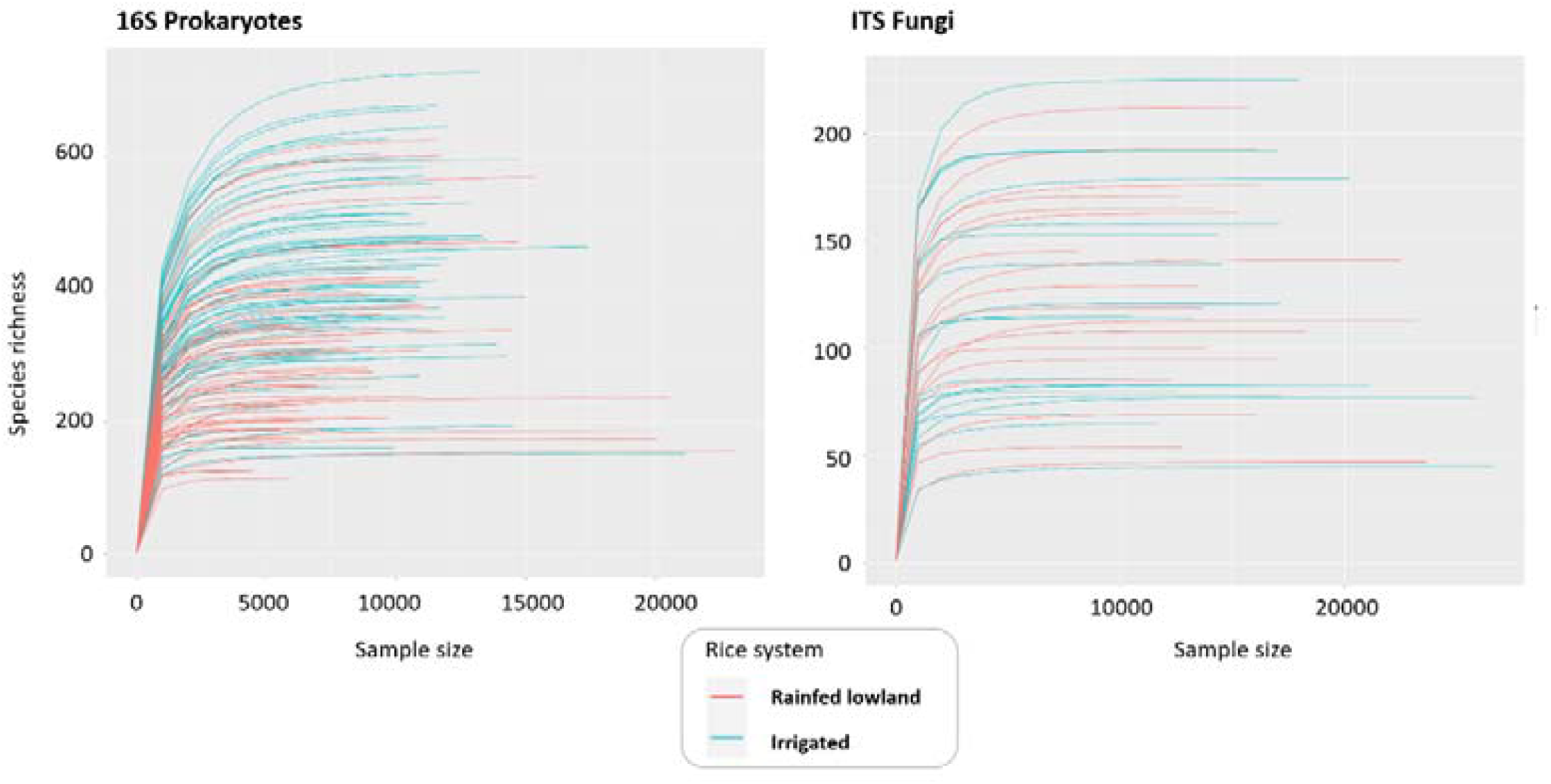
Rarefaction curves (plot of the number of ASVs obtained against the number of analysed reads) for each analyzed samples. On the left, are shown 16S metabarcoding data, representing prokaryotic communities. On the right are presented the ITS metabarcoding data, representing fungal communities.

**Figure S3:**
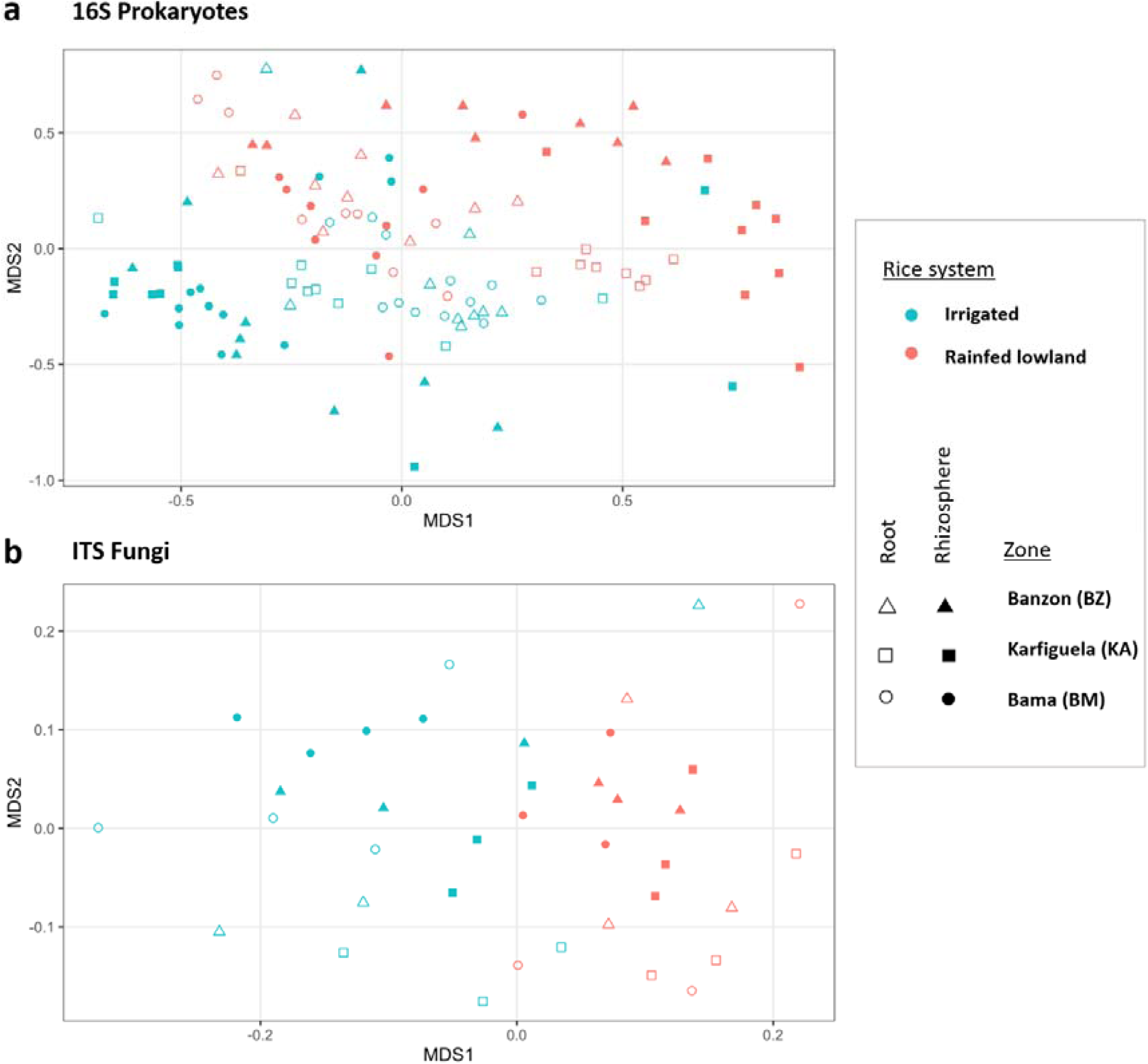
NMDS ordination showing the three factors identified as drivers of the structuration of rice root microbial communities: the color of points represent the rice growing system (irrigated vs rainfed lowland), while the shape shows the compartments (rhizosphere vs roots) and the geographical zone (Banzon, Karfiguela and Bama). a. Analysis based on 16S rRNA gene reflecting Prokaryote communities. One point corresponds to one plant. b. Analysis based on ITS reflecting fungal communities. One point corresponds to one field.

**Figure S4:**
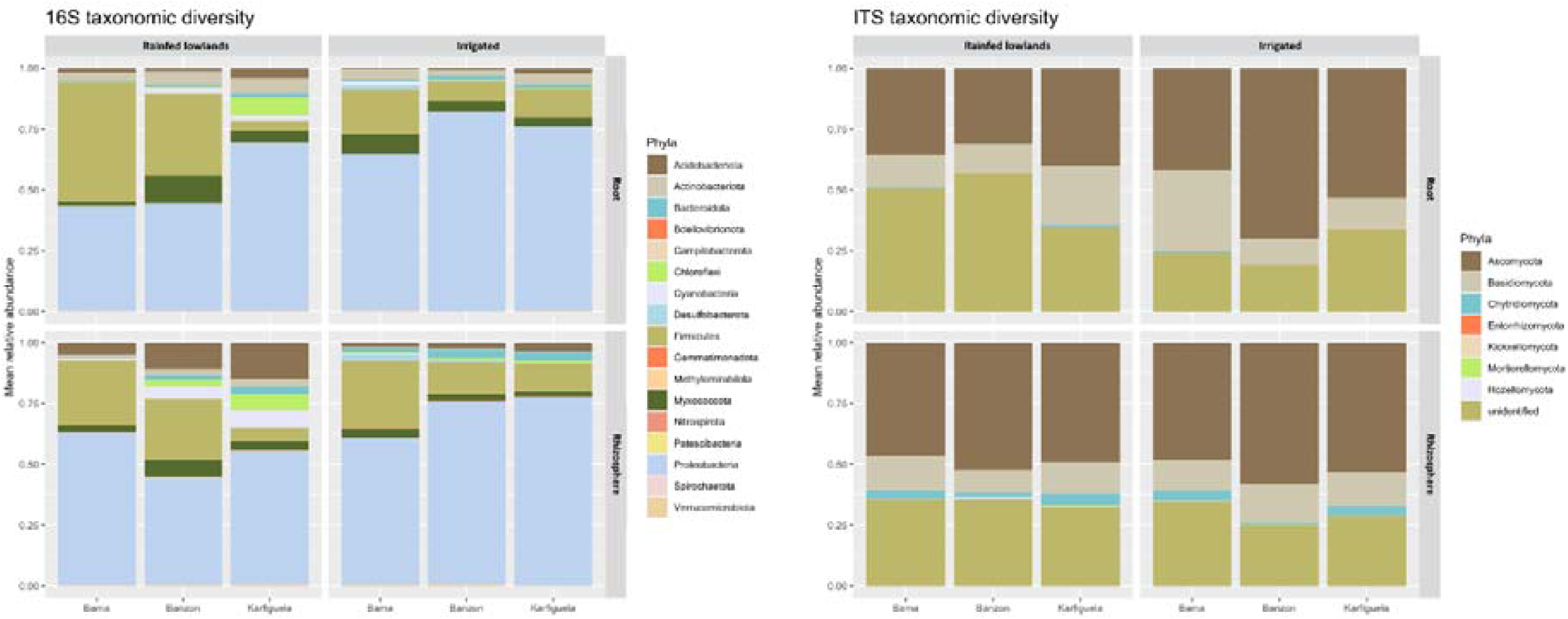
Prokaryote (16S) and fungi (ITS) taxonomic diversity obtained for each study site and each compartment

**Figure S5:**
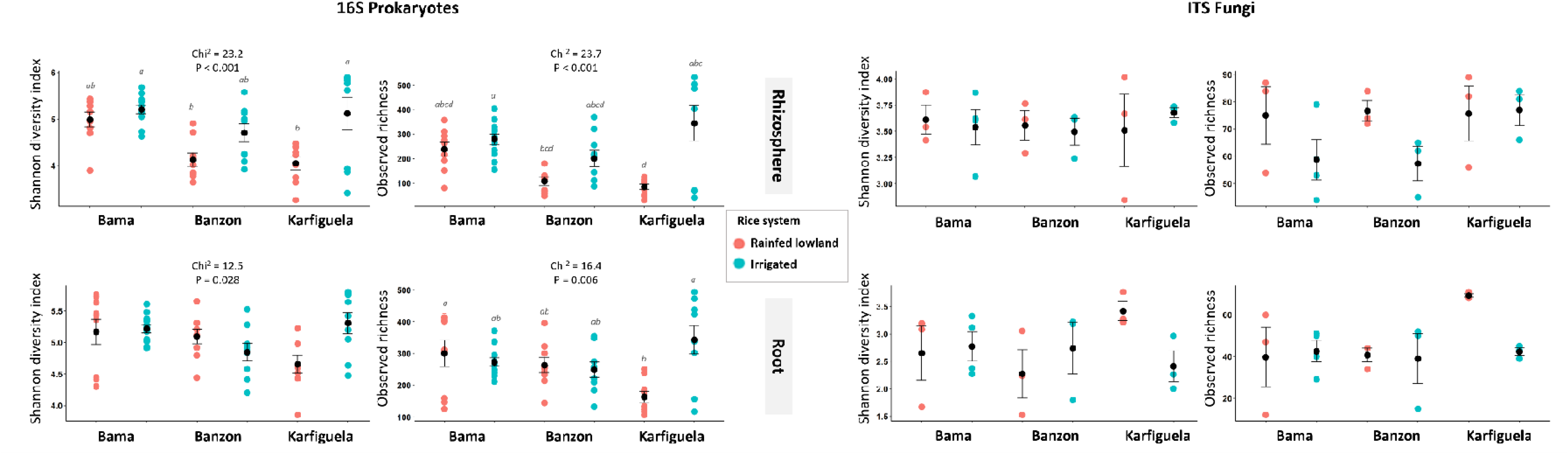
Comparison of root-associated microbiota α-diversity in the six study sites: for Prokaryotes (16S data) on the left and for fungi (ITS data) on the right. Data obtained for the rhizosphere compartment are presented on top and roots data are on the bottom of the figure. The study sites from irrigated areas are represented in blue, while the ones from rainfed lowlands appears in red.

**Table S1.**
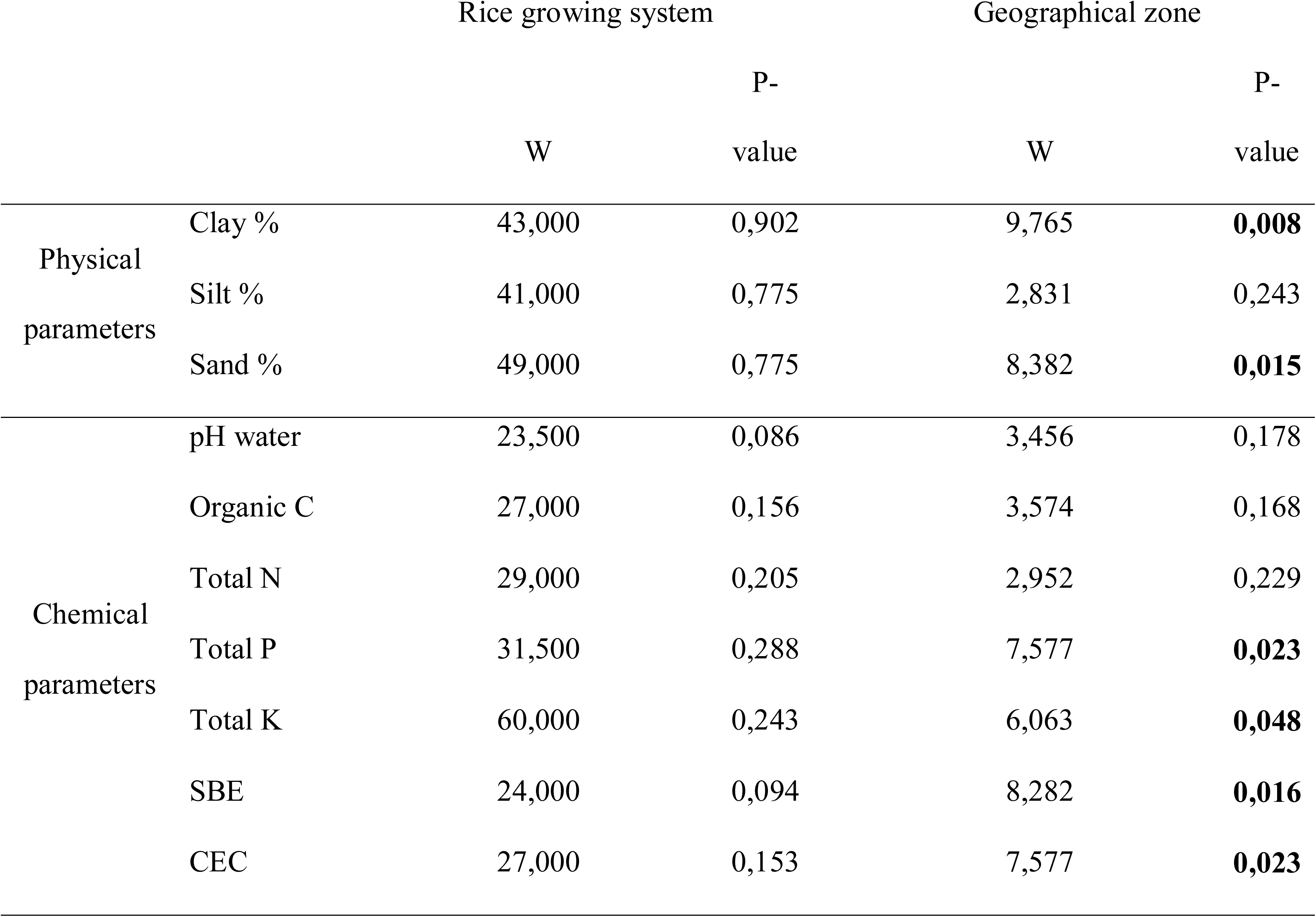
Non-parametric tests (Wilcoxon tests) on the soil physico-chemical parameters

**Table S2.**
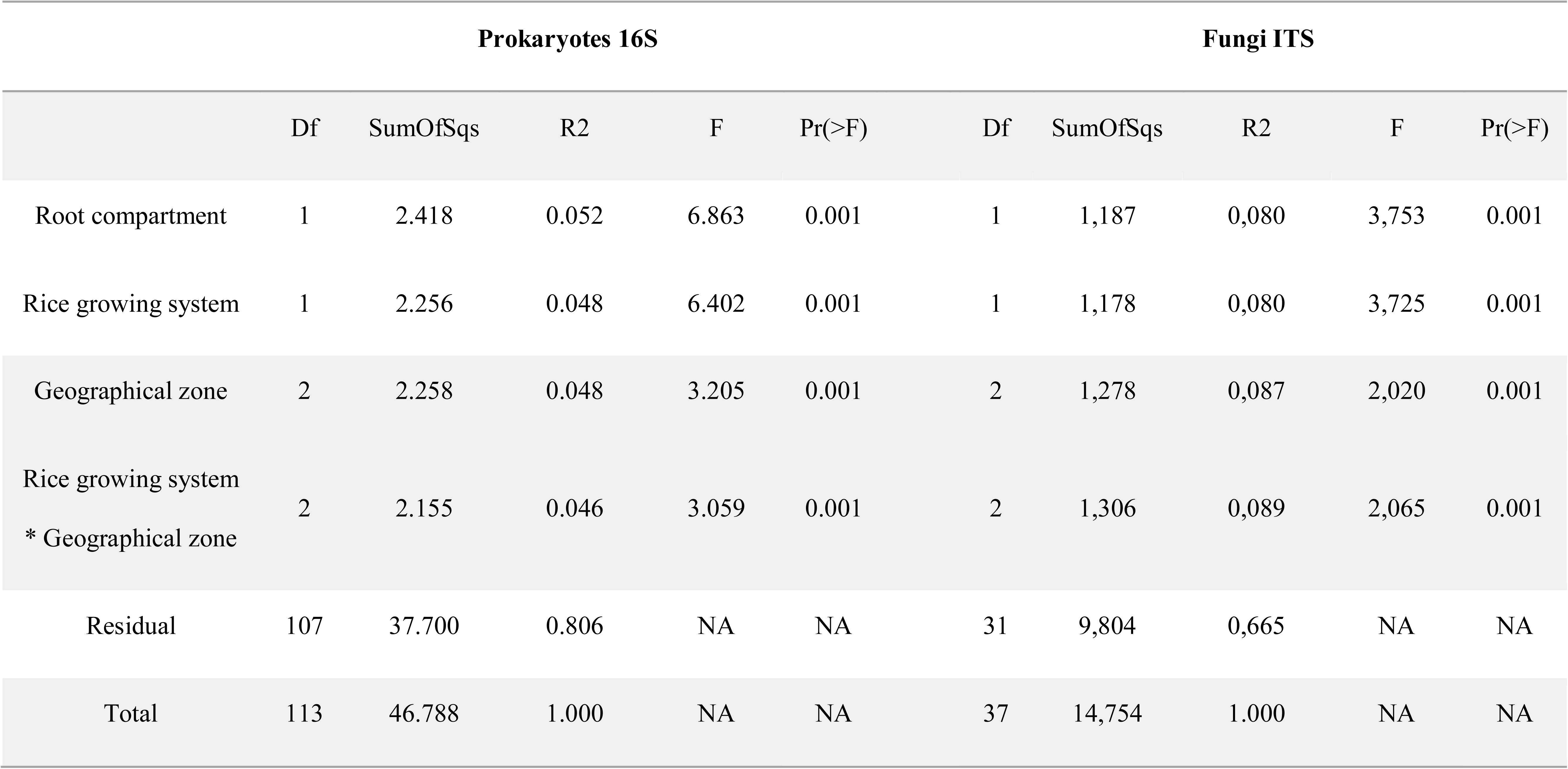
Results of permanova analysis on 16S and ITS microbiome data (all dataset for each of the two markers)

**Table S3.**
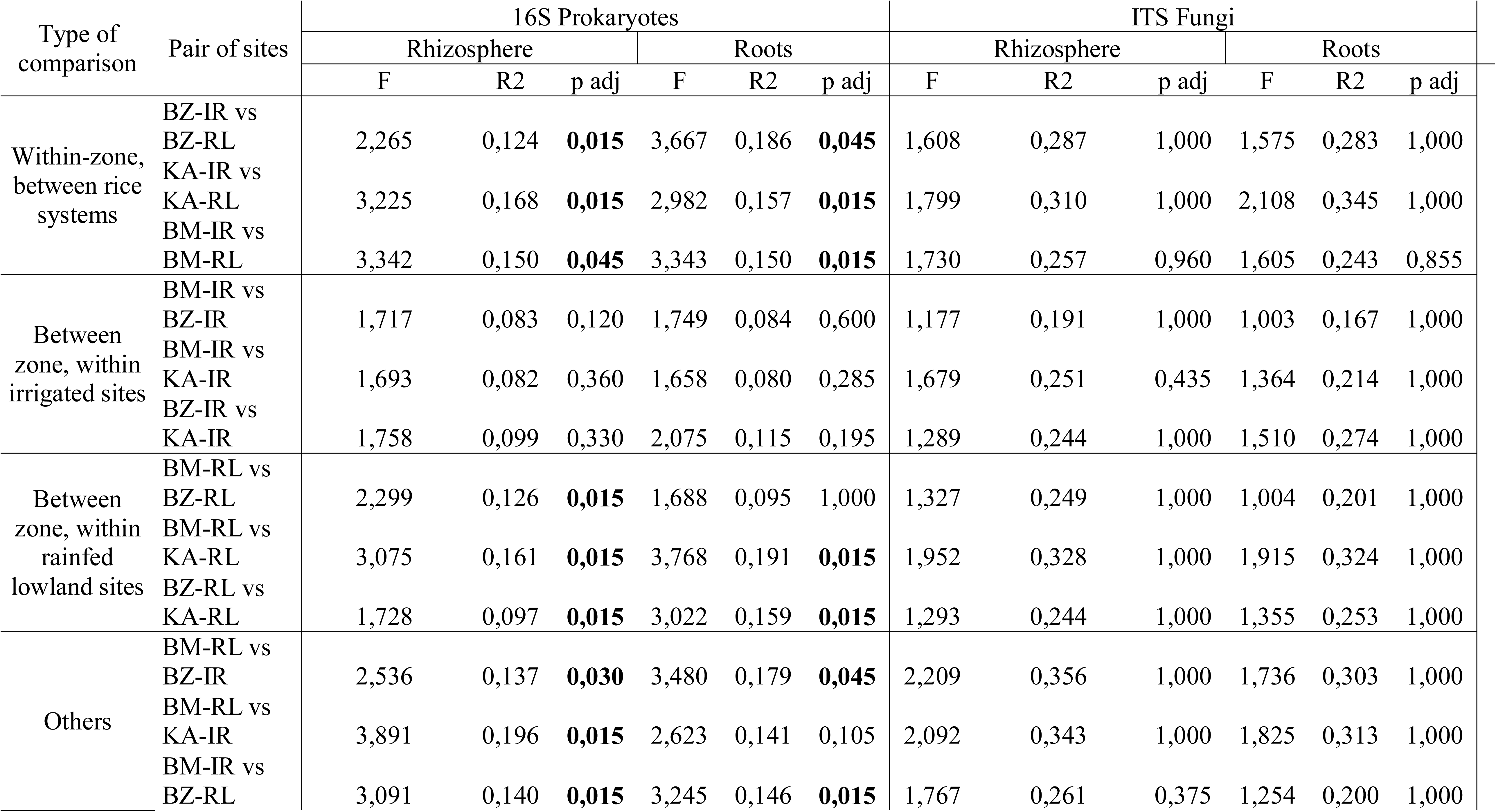
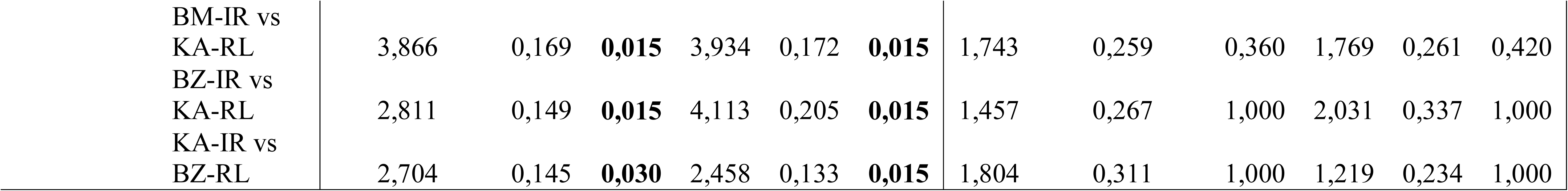
Results of Posthoc tests realized after permanova analysis on 16S and ITS microbiome data, and each compartment independently.

**Table S4.**
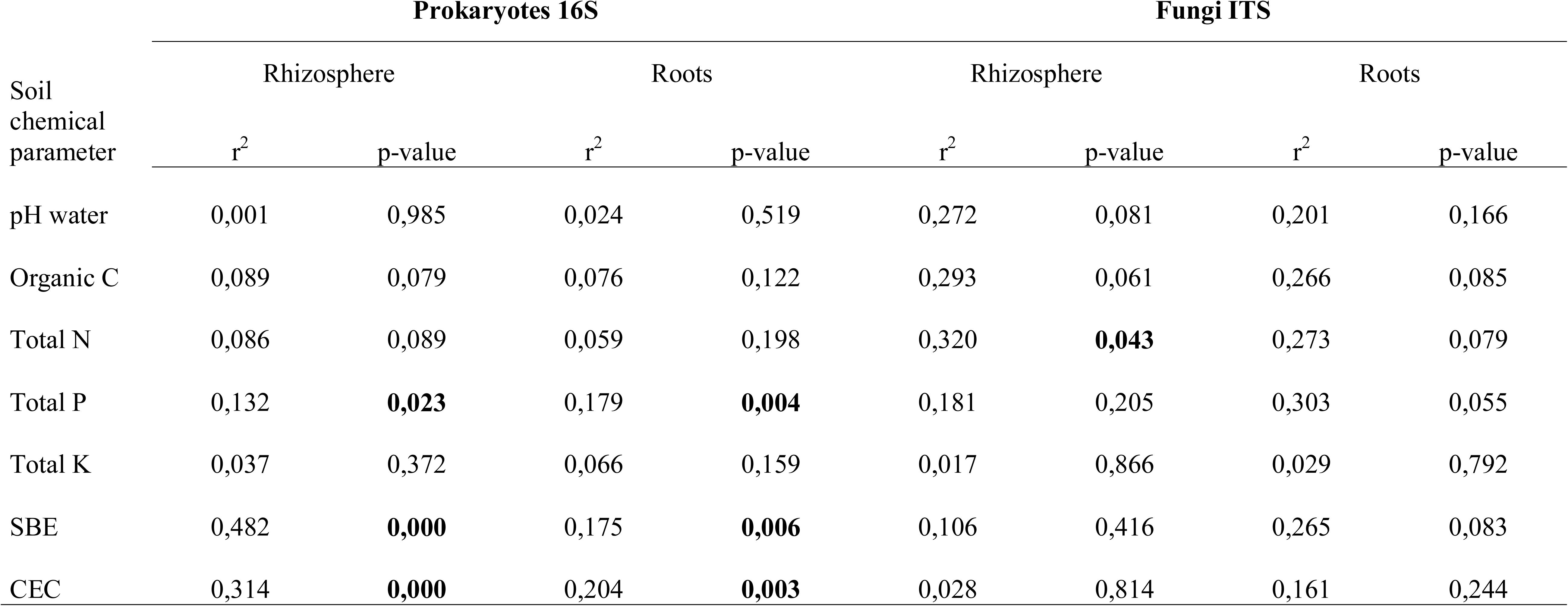
Results of the statistical analyses testing for the effect of soil chemical parameters on microbiome communities (each compartment analyzed separately).

**Table S5.**
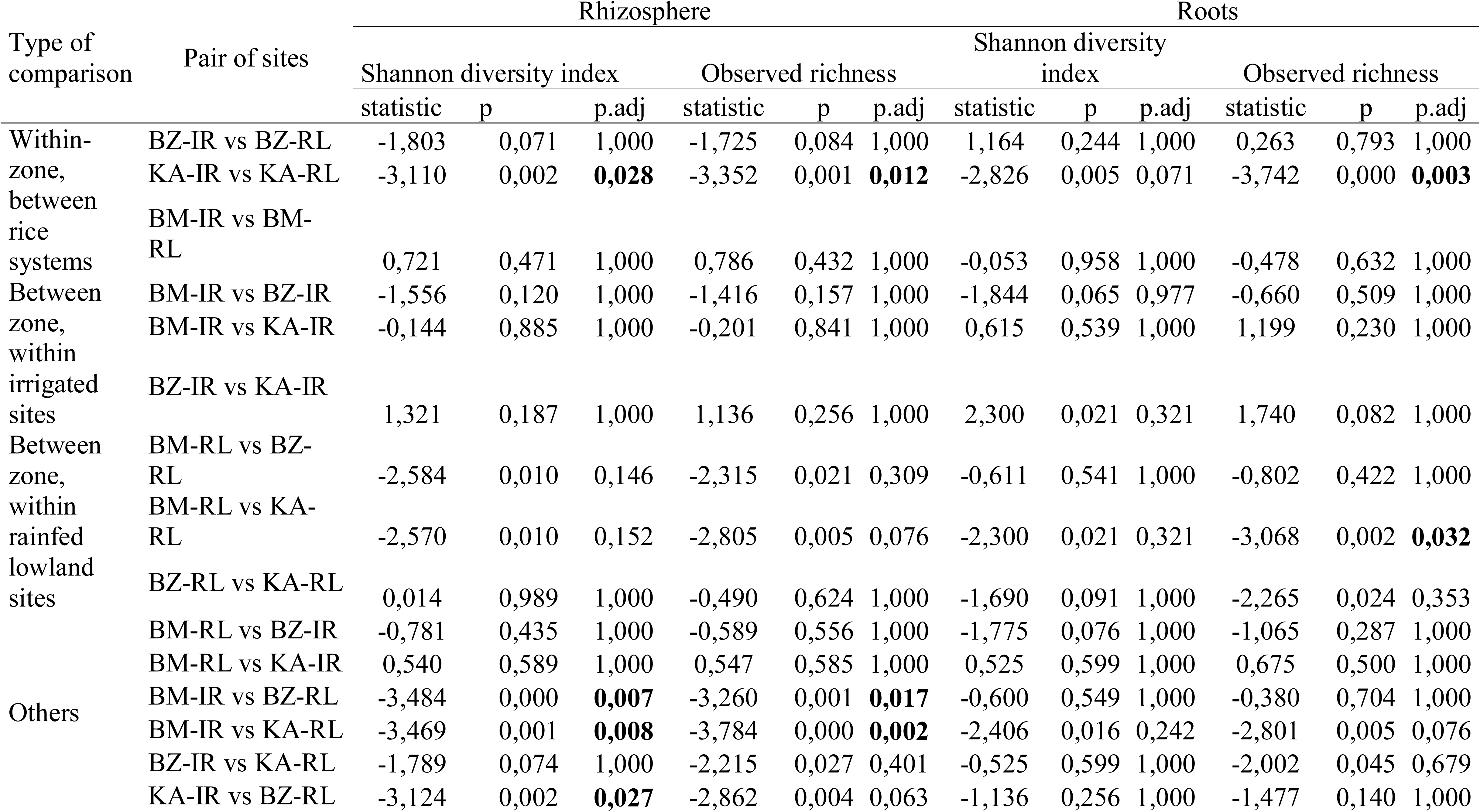
Results of Posthoc tests on the effect of the particular site on alpha diversity indices (Shannon diversity index and observed richness) for 16S microbiome data only and for each compartment independently.

**Table S6.**
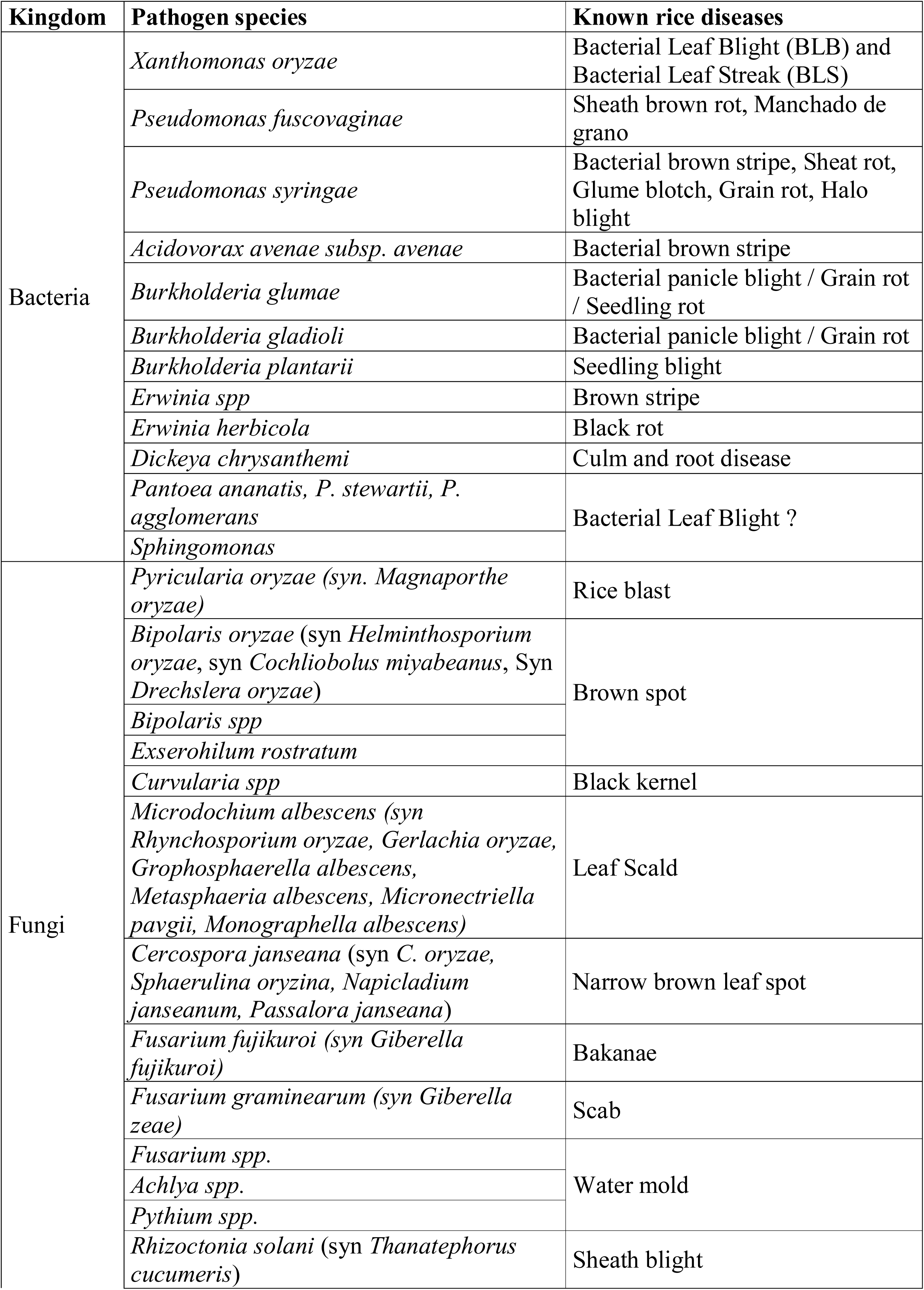
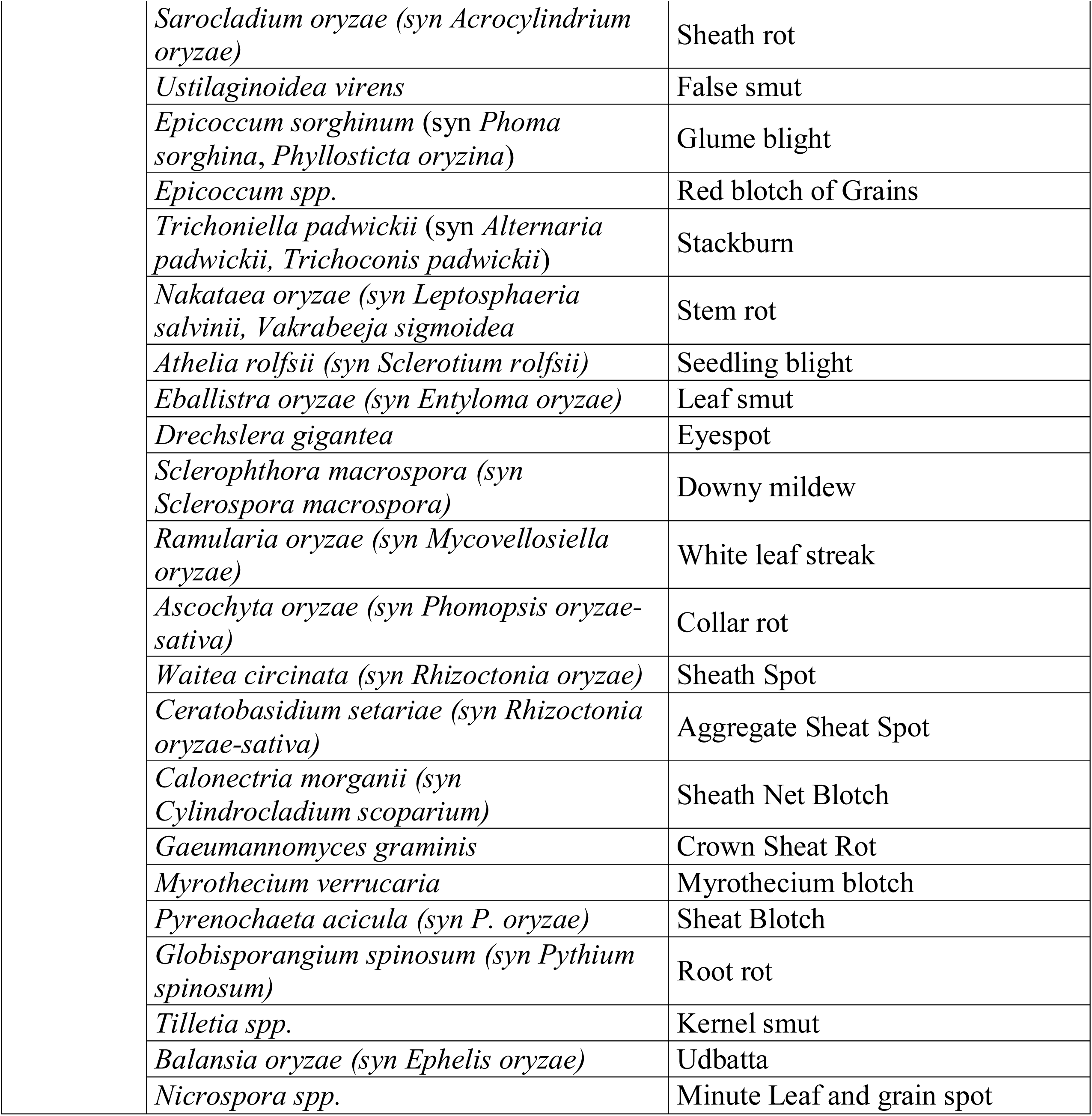
List of the pathogen species searched for within the microbiome data

