## Supplementary files for "The impact of the rice production system (irrigated vs lowland) on root-associated microbiome from farmer’s fields in western Burkina Faso"

**Figure S1:** Monthly precipitation (on the left) and average temperature (on the right) over 1970-2000 period, for each of the six study sites in Western Burkina Faso during the rice growing season (July to November).

WorldClim 2 data (Fick & Hijmans, 2017): <https://worldclim.org/data/worldclim21.html>

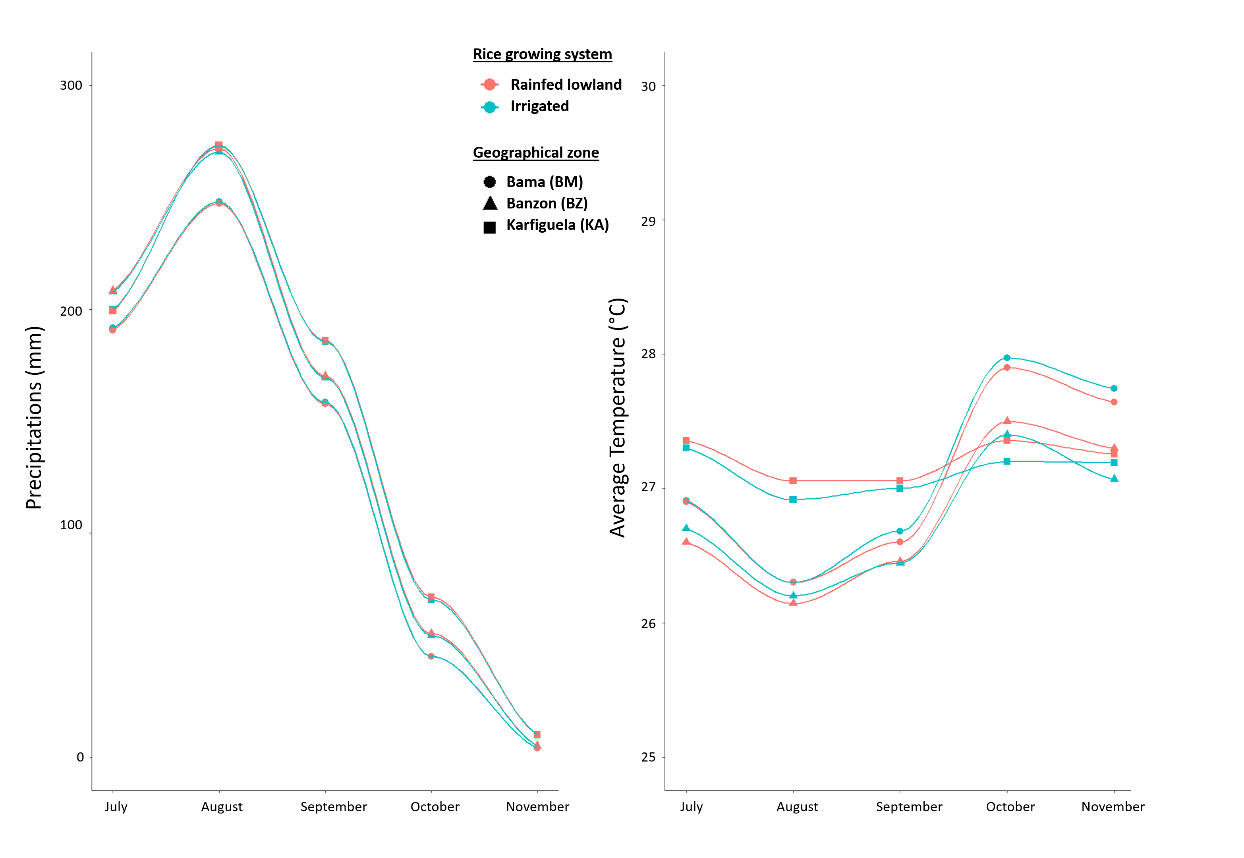

**Figure S2:** Rarefaction curves (plot of the number of ASVs obtained against the number of analysed reads) for each analyzed samples. On the left, are shown 16S metabarcoding data, representing prokaryotic communities. On the right are presented the ITS metabarcoding data, representing fungal communities.

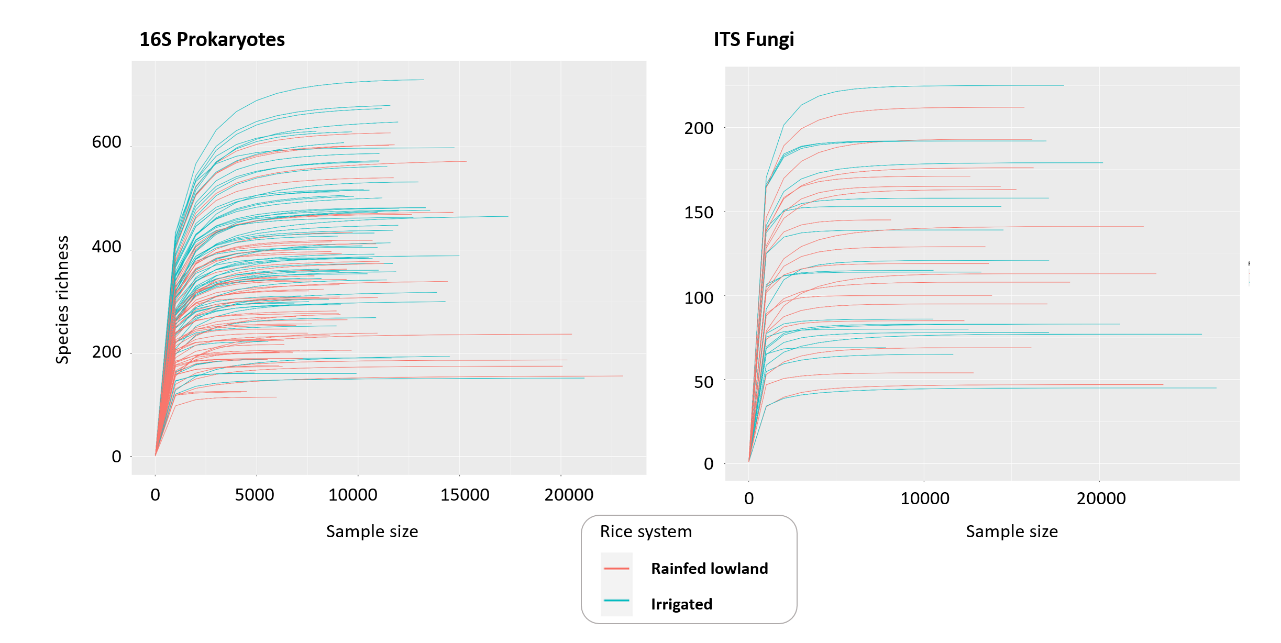

**Figure S3:** NMDS ordination showing the three factors identified as drivers of the structuration of rice root microbial communities: the color of points represent the rice growing system (irrigated vs rainfed lowland), while the shape shows the compartments (rhizosphere vs roots) and the geographical zone (Banzon, Karfiguela and Bama).

a. Analysis based on 16S rRNA gene reflecting Prokaryote communities. One point corresponds to one plant.

b. Analysis based on ITS reflecting fungal communities. One point corresponds to one field.

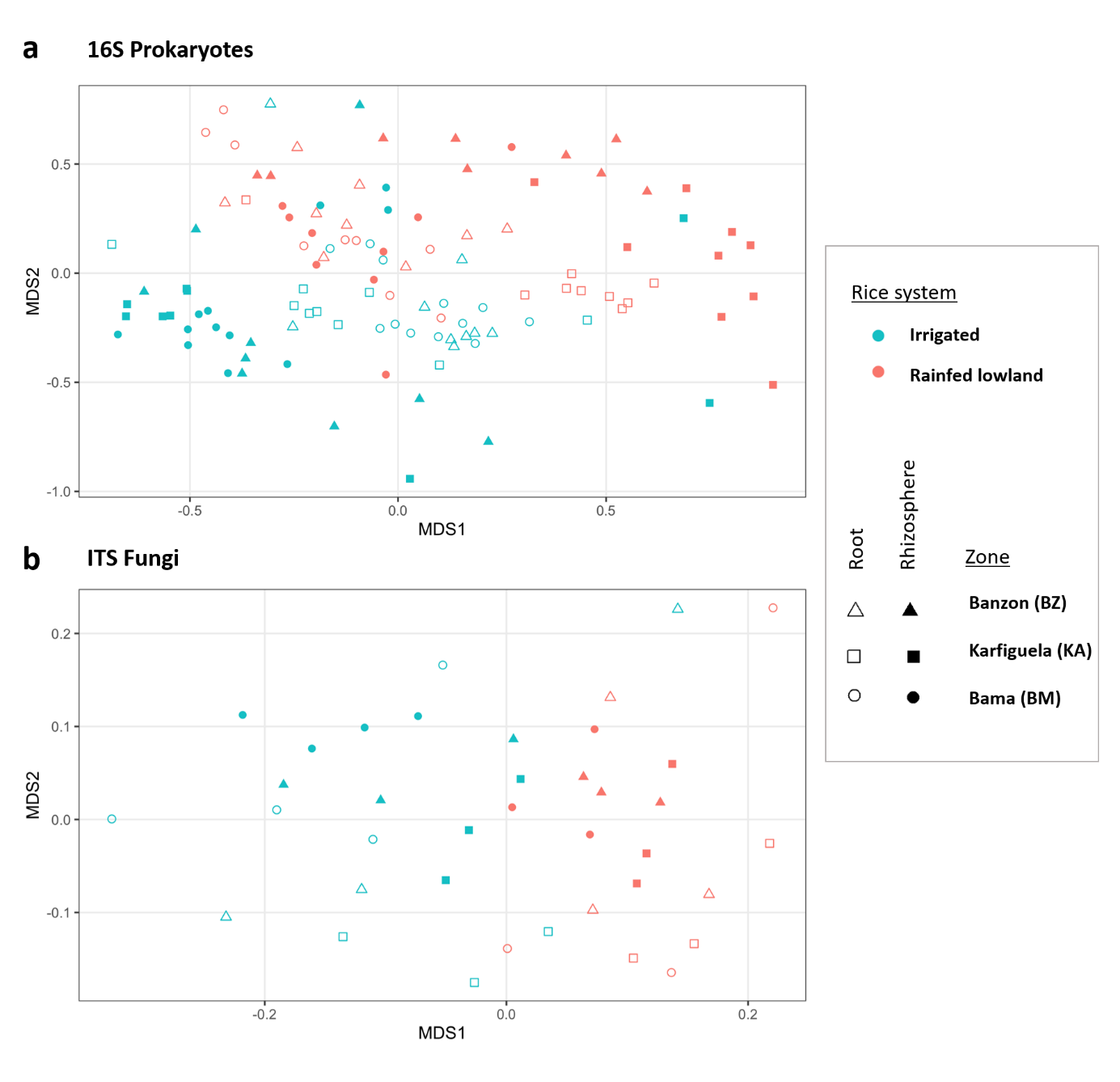

**Figure S4:** Prokaryote (16S) and fungi (ITS) taxonomic diversity obtained for each study site and each compartment

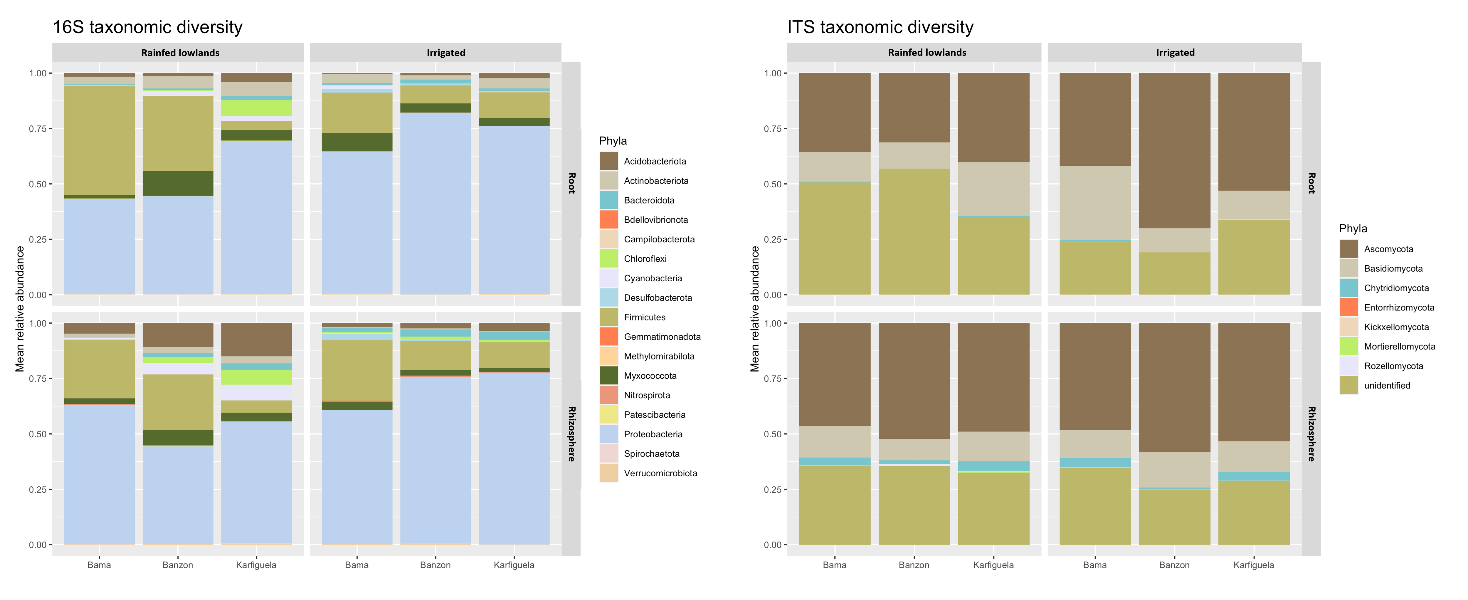

**Figure S5:** Comparison of root-associated microbiota α-diversity in the six study sites: for Prokaryotes (16S data) on the left and for fungi (ITS data) on the right.

Data obtained for the rhizosphere compartment are presented on top and roots data are on the bottom of the figure. The study sites from irrigated areas are represented in blue, while the ones from rainfed lowlands appears in red.

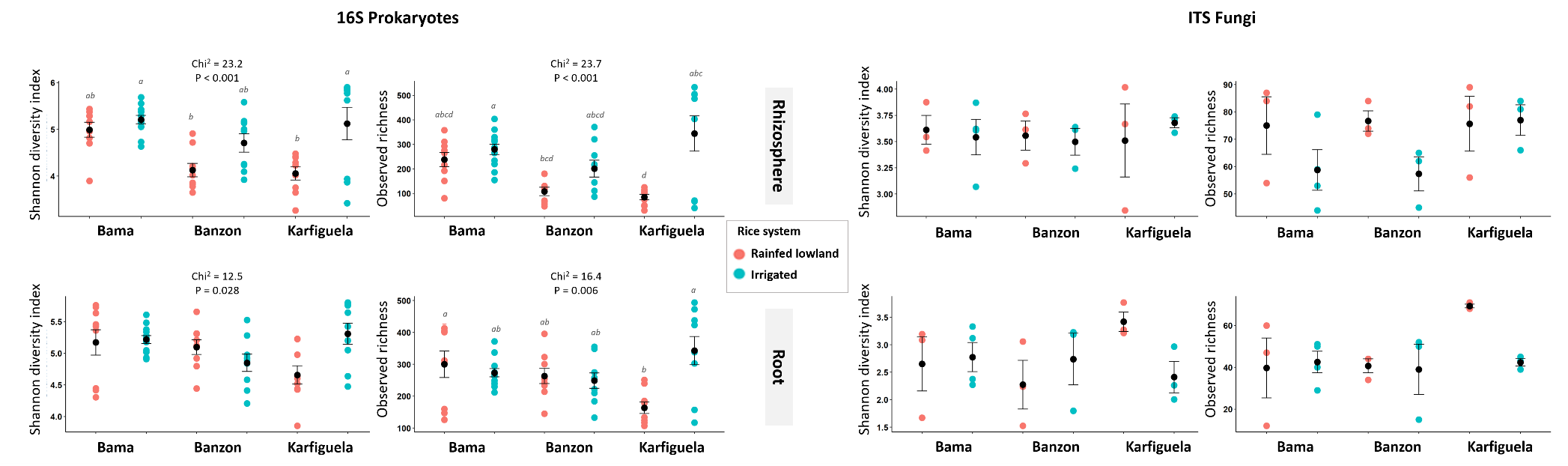

**Table S1**

Non-parametric tests (Wilcoxon tests) on the soil physico-chemical parameters

|  |  | Rice growing system | |  | Geographical zone | |
| --- | --- | --- | --- | --- | --- | --- |
|  |  | W | P-value |  | W | P-value |
| Physical parameters | Clay % | 43,000 | 0,902 |  | 9,765 | **0,008** |
|  | Silt % | 41,000 | 0,775 |  | 2,831 | 0,243 |
|  | Sand % | 49,000 | 0,775 |  | 8,382 | **0,015** |
| Chemical parameters | pH water | 23,500 | 0,086 |  | 3,456 | 0,178 |
|  | Organic C | 27,000 | 0,156 |  | 3,574 | 0,168 |
|  | Total N | 29,000 | 0,205 |  | 2,952 | 0,229 |
|  | Total P | 31,500 | 0,288 |  | 7,577 | **0,023** |
|  | Total K | 60,000 | 0,243 |  | 6,063 | **0,048** |
|  | SBE | 24,000 | 0,094 |  | 8,282 | **0,016** |
|  | CEC | 27,000 | 0,153 |  | 7,577 | **0,023** |

**Table S2**

Results of permanova analysis on 16S and ITS microbiome data (all dataset for each of the two markers)

|  | **Prokaryotes 16S** | | | | |  | | **Fungi ITS** | | | | | |
| --- | --- | --- | --- | --- | --- | --- | --- | --- | --- | --- | --- | --- | --- |
|  | Df | SumOfSqs | R2 | F | Pr(>F) | |  | | Df | SumOfSqs | R2 | F | Pr(>F) |
| Root compartment | 1 | 2.418 | 0.052 | 6.863 | 0.001 | |  | | 1 | 1,187 | 0,080 | 3,753 | 0.001 |
| Rice growing system | 1 | 2.256 | 0.048 | 6.402 | 0.001 | |  | | 1 | 1,178 | 0,080 | 3,725 | 0.001 |
| Geographical zone | 2 | 2.258 | 0.048 | 3.205 | 0.001 | |  | | 2 | 1,278 | 0,087 | 2,020 | 0.001 |
| Rice growing system  * Geographical zone | 2 | 2.155 | 0.046 | 3.059 | 0.001 | |  | | 2 | 1,306 | 0,089 | 2,065 | 0.001 |
| Residual | 107 | 37.700 | 0.806 | NA | NA | |  | | 31 | 9,804 | 0,665 | NA | NA |
| Total | 113 | 46.788 | 1.000 | NA | NA | |  | | 37 | 14,754 | 1.000 | NA | NA |

**Table S3**

Results of Posthoc tests realized after permanova analysis on 16S and ITS microbiome data, and each compartment independently.

| Type of comparison | Pair of sites | 16S Prokaryotes | | | | | | ITS Fungi | | | | | |
| --- | --- | --- | --- | --- | --- | --- | --- | --- | --- | --- | --- | --- | --- |
|  |  | Rhizosphere | | | Roots | | | Rhizosphere | | | Roots | | |
|  |  | F | R2 | p adj | F | R2 | p adj | F | R2 | p adj | F | R2 | p adj |
| Within-zone, between rice systems | BZ-IR vs BZ-RL | 2,265 | 0,124 | **0,015** | 3,667 | 0,186 | **0,045** | 1,608 | 0,287 | 1,000 | 1,575 | 0,283 | 1,000 |
|  | KA-IR vs KA-RL | 3,225 | 0,168 | **0,015** | 2,982 | 0,157 | **0,015** | 1,799 | 0,310 | 1,000 | 2,108 | 0,345 | 1,000 |
|  | BM-IR vs BM-RL | 3,342 | 0,150 | **0,045** | 3,343 | 0,150 | **0,015** | 1,730 | 0,257 | 0,960 | 1,605 | 0,243 | 0,855 |
| Between zone, within irrigated sites | BM-IR vs BZ-IR | 1,717 | 0,083 | 0,120 | 1,749 | 0,084 | 0,600 | 1,177 | 0,191 | 1,000 | 1,003 | 0,167 | 1,000 |
|  | BM-IR vs KA-IR | 1,693 | 0,082 | 0,360 | 1,658 | 0,080 | 0,285 | 1,679 | 0,251 | 0,435 | 1,364 | 0,214 | 1,000 |
|  | BZ-IR vs KA-IR | 1,758 | 0,099 | 0,330 | 2,075 | 0,115 | 0,195 | 1,289 | 0,244 | 1,000 | 1,510 | 0,274 | 1,000 |
| Between zone, within rainfed lowland sites | BM-RL vs BZ-RL | 2,299 | 0,126 | **0,015** | 1,688 | 0,095 | 1,000 | 1,327 | 0,249 | 1,000 | 1,004 | 0,201 | 1,000 |
|  | BM-RL vs KA-RL | 3,075 | 0,161 | **0,015** | 3,768 | 0,191 | **0,015** | 1,952 | 0,328 | 1,000 | 1,915 | 0,324 | 1,000 |
|  | BZ-RL vs KA-RL | 1,728 | 0,097 | **0,015** | 3,022 | 0,159 | **0,015** | 1,293 | 0,244 | 1,000 | 1,355 | 0,253 | 1,000 |
| Others | BM-RL vs BZ-IR | 2,536 | 0,137 | **0,030** | 3,480 | 0,179 | **0,045** | 2,209 | 0,356 | 1,000 | 1,736 | 0,303 | 1,000 |
|  | BM-RL vs KA-IR | 3,891 | 0,196 | **0,015** | 2,623 | 0,141 | 0,105 | 2,092 | 0,343 | 1,000 | 1,825 | 0,313 | 1,000 |
|  | BM-IR vs BZ-RL | 3,091 | 0,140 | **0,015** | 3,245 | 0,146 | **0,015** | 1,767 | 0,261 | 0,375 | 1,254 | 0,200 | 1,000 |
|  | BM-IR vs KA-RL | 3,866 | 0,169 | **0,015** | 3,934 | 0,172 | **0,015** | 1,743 | 0,259 | 0,360 | 1,769 | 0,261 | 0,420 |
|  | BZ-IR vs KA-RL | 2,811 | 0,149 | **0,015** | 4,113 | 0,205 | **0,015** | 1,457 | 0,267 | 1,000 | 2,031 | 0,337 | 1,000 |
|  | KA-IR vs BZ-RL | 2,704 | 0,145 | **0,030** | 2,458 | 0,133 | **0,015** | 1,804 | 0,311 | 1,000 | 1,219 | 0,234 | 1,000 |

**Table S4**

Results of the statistical analyses testing for the effect of soil chemical parameters on microbiome communities (each compartment analyzed separately).

|  | **Prokaryotes 16S** | | | | **Fungi ITS** | | | |
| --- | --- | --- | --- | --- | --- | --- | --- | --- |
| Soil chemical parameter | Rhizosphere | | Roots | | Rhizosphere | | Roots | |
|  | r^2^ | p-value | r^2^ | p-value | r^2^ | p-value | r^2^ | p-value |
| pH water | 0,001 | 0,985 | 0,024 | 0,519 | 0,272 | 0,081 | 0,201 | 0,166 |
| Organic C | 0,089 | 0,079 | 0,076 | 0,122 | 0,293 | 0,061 | 0,266 | 0,085 |
| Total N | 0,086 | 0,089 | 0,059 | 0,198 | 0,320 | **0,043** | 0,273 | 0,079 |
| Total P | 0,132 | **0,023** | 0,179 | **0,004** | 0,181 | 0,205 | 0,303 | 0,055 |
| Total K | 0,037 | 0,372 | 0,066 | 0,159 | 0,017 | 0,866 | 0,029 | 0,792 |
| SBE | 0,482 | **0,000** | 0,175 | **0,006** | 0,106 | 0,416 | 0,265 | 0,083 |
| CEC | 0,314 | **0,000** | 0,204 | **0,003** | 0,028 | 0,814 | 0,161 | 0,244 |

**Table S5** Results of Posthoc tests on the effect of the particular site on alpha diversity indices (Shannon diversity index and observed richness) for 16S microbiome data only and for each compartment independently.

| Type of comparison | Pair of sites | Rhizosphere | | | | | | Roots | | | | | |
| --- | --- | --- | --- | --- | --- | --- | --- | --- | --- | --- | --- | --- | --- |
|  |  | Shannon diversity index | | | Observed richness | | | Shannon diversity index | | | Observed richness | | |
|  |  | statistic | p | p.adj | statistic | p | p.adj | statistic | p | p.adj | statistic | p | p.adj |
| Within-zone, between rice systems | BZ-IR vs BZ-RL | -1,803 | 0,071 | 1,000 | -1,725 | 0,084 | 1,000 | 1,164 | 0,244 | 1,000 | 0,263 | 0,793 | 1,000 |
|  | KA-IR vs KA-RL | -3,110 | 0,002 | **0,028** | -3,352 | 0,001 | **0,012** | -2,826 | 0,005 | 0,071 | -3,742 | 0,000 | **0,003** |
|  | BM-IR vs BM-RL | 0,721 | 0,471 | 1,000 | 0,786 | 0,432 | 1,000 | -0,053 | 0,958 | 1,000 | -0,478 | 0,632 | 1,000 |
| Between zone, within irrigated sites | BM-IR vs BZ-IR | -1,556 | 0,120 | 1,000 | -1,416 | 0,157 | 1,000 | -1,844 | 0,065 | 0,977 | -0,660 | 0,509 | 1,000 |
|  | BM-IR vs KA-IR | -0,144 | 0,885 | 1,000 | -0,201 | 0,841 | 1,000 | 0,615 | 0,539 | 1,000 | 1,199 | 0,230 | 1,000 |
|  | BZ-IR vs KA-IR | 1,321 | 0,187 | 1,000 | 1,136 | 0,256 | 1,000 | 2,300 | 0,021 | 0,321 | 1,740 | 0,082 | 1,000 |
| Between zone, within rainfed lowland sites | BM-RL vs BZ-RL | -2,584 | 0,010 | 0,146 | -2,315 | 0,021 | 0,309 | -0,611 | 0,541 | 1,000 | -0,802 | 0,422 | 1,000 |
|  | BM-RL vs KA-RL | -2,570 | 0,010 | 0,152 | -2,805 | 0,005 | 0,076 | -2,300 | 0,021 | 0,321 | -3,068 | 0,002 | **0,032** |
|  | BZ-RL vs KA-RL | 0,014 | 0,989 | 1,000 | -0,490 | 0,624 | 1,000 | -1,690 | 0,091 | 1,000 | -2,265 | 0,024 | 0,353 |
| Others | BM-RL vs BZ-IR | -0,781 | 0,435 | 1,000 | -0,589 | 0,556 | 1,000 | -1,775 | 0,076 | 1,000 | -1,065 | 0,287 | 1,000 |
|  | BM-RL vs KA-IR | 0,540 | 0,589 | 1,000 | 0,547 | 0,585 | 1,000 | 0,525 | 0,599 | 1,000 | 0,675 | 0,500 | 1,000 |
|  | BM-IR vs BZ-RL | -3,484 | 0,000 | **0,007** | -3,260 | 0,001 | **0,017** | -0,600 | 0,549 | 1,000 | -0,380 | 0,704 | 1,000 |
|  | BM-IR vs KA-RL | -3,469 | 0,001 | **0,008** | -3,784 | 0,000 | **0,002** | -2,406 | 0,016 | 0,242 | -2,801 | 0,005 | 0,076 |
|  | BZ-IR vs KA-RL | -1,789 | 0,074 | 1,000 | -2,215 | 0,027 | 0,401 | -0,525 | 0,599 | 1,000 | -2,002 | 0,045 | 0,679 |
|  | KA-IR vs BZ-RL | -3,124 | 0,002 | **0,027** | -2,862 | 0,004 | 0,063 | -1,136 | 0,256 | 1,000 | -1,477 | 0,140 | 1,000 |

**Table S6**

List of the pathogen species searched for within the microbiome data

| **Kingdom** | **Pathogen species** | **Known rice diseases** |
| --- | --- | --- |
| Bacteria | *Xanthomonas oryzae* | Bacterial Leaf Blight (BLB) and Bacterial Leaf Streak (BLS) |
|  | *Pseudomonas fuscovaginae* | Sheath brown rot, Manchado de grano |
|  | *Pseudomonas syringae* | Bacterial brown stripe, Sheat rot, Glume blotch, Grain rot, Halo blight |
|  | *Acidovorax avenae subsp. avenae* | Bacterial brown stripe |
|  | *Burkholderia glumae* | Bacterial panicle blight / Grain rot / Seedling rot |
|  | *Burkholderia gladioli* | Bacterial panicle blight / Grain rot |
|  | *Burkholderia plantarii* | Seedling blight |
|  | *Erwinia spp* | Brown stripe |
|  | *Erwinia herbicola* | Black rot |
|  | *Dickeya chrysanthemi* | Culm and root disease |
|  | *Pantoea ananatis, P. stewartii, P. agglomerans* | Bacterial Leaf Blight ? |
|  | *Sphingomonas* |  |
| Fungi | *Pyricularia oryzae (syn. Magnaporthe oryzae)* | Rice blast |
|  | *Bipolaris oryzae* (syn *Helminthosporium oryzae*, syn *Cochliobolus miyabeanus*, Syn *Drechslera oryzae*) | Brown spot |
|  | *Bipolaris spp* |  |
|  | *Exserohilum rostratum* |  |
|  | *Curvularia spp* | Black kernel |
|  | *Microdochium albescens (syn Rhynchosporium oryzae, Gerlachia oryzae, Grophosphaerella albescens, Metasphaeria albescens, Micronectriella pavgii, Monographella albescens)* | Leaf Scald |
|  | *Cercospora janseana* (syn *C. oryzae, Sphaerulina oryzina, Napicladium janseanum, Passalora janseana*) | Narrow brown leaf spot |
|  | *Fusarium fujikuroi (syn Giberella fujikuroi)* | Bakanae |
|  | *Fusarium graminearum (syn Giberella zeae)* | Scab |
|  | *Fusarium spp.* | Water mold |
|  | *Achlya spp.* |  |
|  | *Pythium spp.* |  |
|  | *Rhizoctonia solani* (syn *Thanatephorus cucumeris*) | Sheath blight |
|  | *Sarocladium oryzae (syn Acrocylindrium oryzae)* | Sheath rot |
|  | *Ustilaginoidea virens* | False smut |
|  | *Epicoccum sorghinum* (syn *Phoma sorghina*, *Phyllosticta oryzina*) | Glume blight |
|  | *Epicoccum spp.* | Red blotch of Grains |
|  | *Trichoniella padwickii* (syn *Alternaria padwickii, Trichoconis padwickii*) | Stackburn |
|  | *Nakataea oryzae (syn Leptosphaeria salvinii, Vakrabeeja sigmoidea* | Stem rot |
|  | *Athelia rolfsii (syn Sclerotium rolfsii)* | Seedling blight |
|  | *Eballistra oryzae (syn Entyloma oryzae)* | Leaf smut |
|  | *Drechslera gigantea* | Eyespot |
|  | *Sclerophthora macrospora (syn Sclerospora macrospora)* | Downy mildew |
|  | *Ramularia oryzae (syn Mycovellosiella oryzae)* | White leaf streak |
|  | *Ascochyta oryzae (syn Phomopsis oryzae-sativa)* | Collar rot |
|  | *Waitea circinata (syn Rhizoctonia oryzae)* | Sheath Spot |
|  | *Ceratobasidium setariae (syn Rhizoctonia oryzae-sativa)* | Aggregate Sheat Spot |
|  | *Calonectria morganii (syn Cylindrocladium scoparium)* | Sheath Net Blotch |
|  | *Gaeumannomyces graminis* | Crown Sheat Rot |
|  | *Myrothecium verrucaria* | Myrothecium blotch |
|  | *Pyrenochaeta acicula (syn P. oryzae)* | Sheat Blotch |
|  | *Globisporangium spinosum (syn Pythium spinosum)* | Root rot |
|  | *Tilletia spp.* | Kernel smut |
|  | *Balansia oryzae (syn Ephelis oryzae)* | Udbatta |
|  | *Nicrospora spp.* | Minute Leaf and grain spot |
